# Auto-regulatory J-domain interactions control Hsp70 recruitment to the DnaJB8 chaperone

**DOI:** 10.1101/2020.01.09.899237

**Authors:** Bryan D. Ryder, Irina Matlahov, Sofia Bali, Jaime Vaquer-Alicea, Patrick C.A. van der Wel, Lukasz A. Joachimiak

## Abstract

The Hsp40/Hsp70 chaperone families combine versatile folding capacity with high substrate specificity, which is mainly facilitated by Hsp40s. The structure and function of many Hsp40s remain poorly understood, particularly oligomeric Hsp40s that suppress protein aggregation. Here, we used a combination of biochemical and structural approaches to shed new light on the domain interactions of the Hsp40 DnaJB8, and how they regulate recruitment of partner Hsp70s. We identify an interaction between the J-Domain (JD) and C-terminal domain (CTD) of DnaJB8 that sequesters the JD surface, preventing Hsp70 interaction. We propose a new model for DnaJB8-Hsp70 regulation, whereby the JD-CTD interaction of DnaJB8 acts as a reversible autoinhibitory switch that can control the binding of Hsp70. These findings suggest that the evolutionarily conserved CTD of DnaJB8 is a regulatory element of chaperone activity in the proteostasis network.

## INTRODUCTION

The cellular chaperone network needs to handle a diversity of protein substrates in numerous different (mis)folded states. This demands a combination of broad versatility and specificity in terms of substrate recognition, even though the central players Hsp70 and Hsp90 are highly conserved. This apparent contradiction is resolved by the Hsp40 (DnaJ) family of proteins, which are chaperones that recruit and regulate the activity of Hsp70 chaperones in refolding misfolded proteins^1-4^. While the human Hsp70 family is highly conserved, the Hsp40 chaperone family encodes 47 diverse members, each with specialized functions in substrate recognition and presumed coordination with Hsp70^5-7^. DnaJ proteins feature a J-domain (JD), which binds to Hsp70s through a conserved electrostatic interaction to trigger ATP hydrolysis by the Hsp70^7-10^. This initiates a conformational rearrangement in the Hsp70 substrate binding domain that helps capture the substrate for folding, refolding, or disaggregation^11^. When misfolded proteins cannot be refolded, some Hsp40s help direct them for degradation^12-13^.

In humans, Hsp40s function as monomers, dimers, or oligomers. Classical Hsp40 members assemble into homo-dimers or hetero-dimers^14^ through conserved C-terminal motifs and bind unfolded substrates through conserved β-barrel C-terminal domains (CTD)^15^. A subset of non-classical Hsp40s, including DnaJB2, DnaJB6b, DnaJB7 and DnaJB8 have a domain architecture that is distinct from the classical dimeric DnaJ orthologs^14-17^. These Hsp40s retain the JD, but have distinct other domains including substantial differences in their CTD structures. Of these, the DnaJB8 and DnaJB6b proteins self-assemble *in vitro* and *in vivo*^16, 18-19^. The role of their CTD remains unclear, as the literature suggests that it either drives oligomerization or mediates intramolecular contacts^16-18, 20^. The oligomers’ structural and dynamic heterogeneity has greatly hindered efforts to study them, yielding for DnaJB6 limited-resolution cryoEM data^19^ or requiring invasive deletion mutations to gain structural insight in soluble mutant variants^16-17^.

Here we examine DnaJB8, which has been shown to be particularly effective at preventing polyglutamine (polyQ) deposition, even more so than the homologous DnaJB6b, despite 63 % sequence identity^16, 18, 20-21^. This indicates that their specific modes of activity are distinct in spite of their similarities in sequence and domain arrangement. Notably, unlike other chaperones that inhibit mutant huntingtin aggregation^22^, DnaJB8 and DnaJB6b are thought to bind directly to polyQ elements and thus are active across the whole family of polyQ diseases^16, 21^. The two proteins have different expression profiles, with DnaJB8 being highly expressed in testes, while DnaJB6b is ubiquitous, which in part explains the deeper knowledge available for the latter protein. While both DnaJB6b and DnaJB8 assemble into soluble oligomers^16, 18-19, 21^, DnaJB8 in particular displays a higher propensity to assemble^16^. Here, we applied a multidisciplinary approach to understand the architecture and dynamics of DnaJB8 in cells and *in vitro*. We used crosslinking mass spectrometry (XL-MS) to identify local intra-domain contacts and long-range contacts. Guided by modeling, we mutated aromatic residues to create a monomeric mutant that maintains the intramolecular domain contacts observed in oligomers in cells and *in vitro*. Solid-state NMR (ssNMR) probed the structural and dynamic order of the solvated oligomers, to reveal dramatic domain-specific differences in (dis)order and a lack of highly flexible regions. Electrostatic interactions control the JD transitioning between an ordered immobilized state and a more mobilized state, which we attribute to JD - CTD interactions that we reconstitute with isolated domains and detect in full-length protein. Finally, we demonstrate that the JD-CTD contacts regulate the recruitment of Hsp70, representing a built-in regulatory mechanism that controls recruitment (and thus activation) of Hsp70.

## RESULTS

### DnaJB8 domain interactions in a cellular context

DnaJB8 encodes three domains C-terminal to the JD (Fig. 1a): a glycine and phenylalanine (G/F) rich domain (Fig. 1a, blue), a serine/threonine rich (S/T) domain (Fig. 1a, cyan) and a C-terminal CTD (Fig. 1a, green). Prior studies have highlighted the ability of DnaJB8 to assemble into oligomers but little is known about DnaJB8 domain interactions in cells^16, 20^. We expressed DnaJB8 fused to a green fluorescent protein (GFP) derivative mClover3 (herein, DnaJB8-Cover) in HEK293 cells (Fig. 1a). DnaJB8-Clover expression leads to the formation of fluorescent juxtanuclear puncta with an approximate maximum diameter of 1.0μm (Fig. 1b) in 39.2±3.1% of the cells (Fig. 1c) while Clover alone expression yielded diffuse fluorescence (Fig. 1b) with little to no puncta (Fig. 1c; 0.44±0.50%). The puncta observed in these cells indicate the presence of ordered aggregates, while the more uniformly dispersed signal is indicative of soluble oligomers and monomers. Decreasing the DnaJB8-Clover expression 3-fold as determined by western blot (Supplementary Fig. 1a) and fluorescence intensity (Supplementary Fig. 1b) yielded only a 2-fold decrease in the number of puncta (16.2±0.08%; Fig. 1c). The frequency of puncta for Clover alone remained below 1% in both experiments (Fig. 1c). Thus, even at reduced levels of expression DnaJB8 can form puncta in cells.

**Figure 1.**
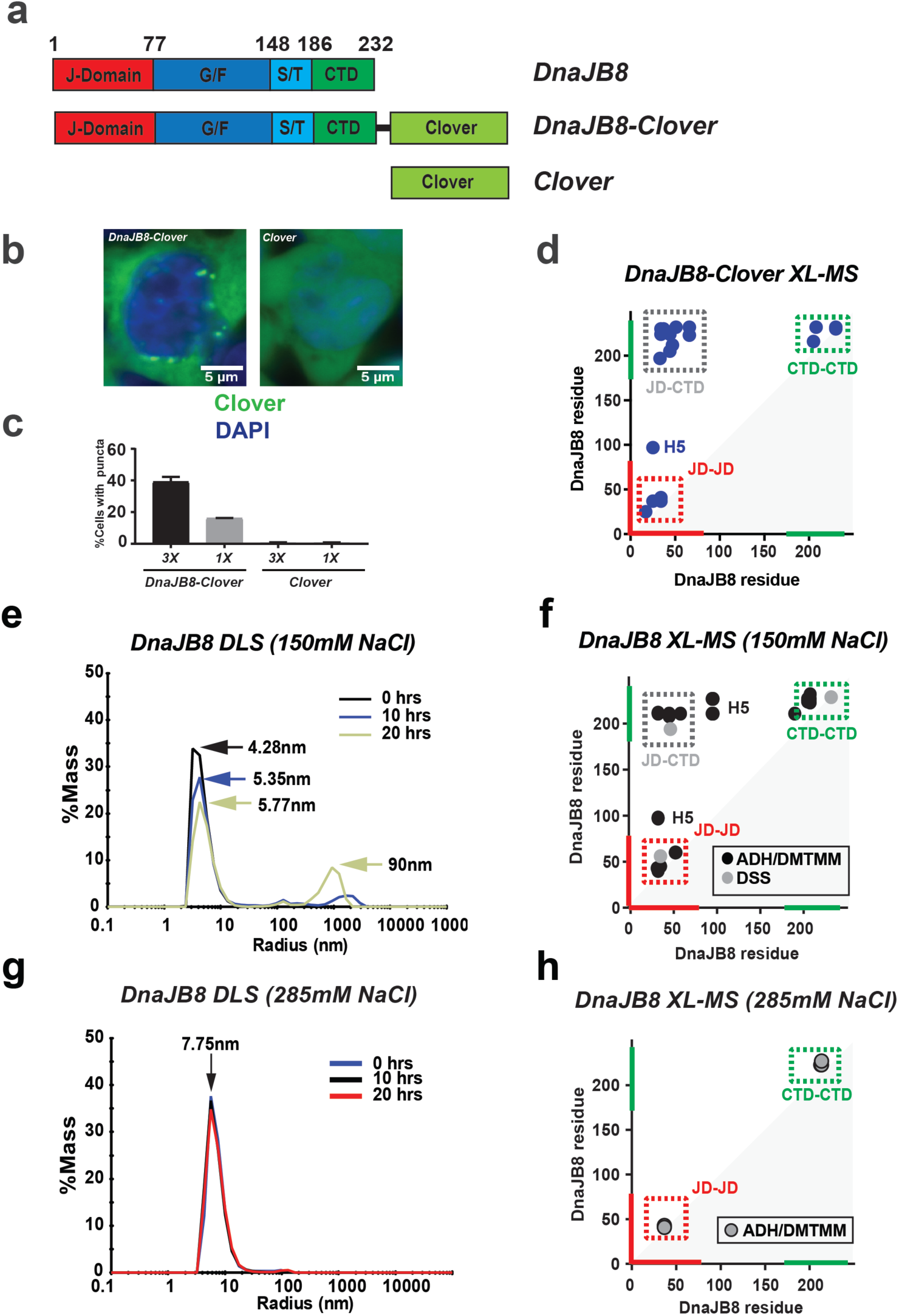
DnaJB8 architecture defined by domain-domain interactions. (**a**) Domain maps for DnaJB8 used in the *in vitro* experiments and the DnaJB-Clover and Clover constructs used in the mammalian cell experiments. DnaJB8 colored according to domain annotation: JD (red), G/F rich (blue), S/T rich (cyan), and CTD (green). Clover is colored pale green. (**b**) Representative images of triplicate populations of 300,000 cells expressing DnaJB8-mClover3(left) and mClover3(right). Clover and DAPI fluorescence signal are shown in green and blue, respectively. 5μm scale bar is shown in white. (**c**) Quantification of DnaJB8-Clover and Clover puncta in high (3X) and low (1X) protein level expressing cell lines. In each analysis at least 2,000 cells were counted by the CellProfiler software. Puncta were manually counted by two independent observers, with data reported as averages with standard deviation. (**d**) XL-MS contact map of DnaJB8-Clover crosslinks identified using DMTMM and ADH. The axes are colored in red and green for JD and CTD, respectively. Crosslink pairs between JD-CTD, JD-JD and CTD-CTD are shown in dashed boxes colored grey, red and green, respectively. Contacts to helix 5 are denoted with H5. (**e)** Histogram of overall R_h_ of DnaJB8 in 1xPBS 150mM NaCl from DLS at times 0hrs (black), 10hrs (blue), and 20hrs (gold) with arrows indicating R_h_ peaks for each time point. Over time, there was a depletion in particle sizes <10nm and an increase in particles ∼100-1000nm. **(f)** XL-MS contact map of DnaJB8 crosslinks identified using DMTMM and ADH (black) and DSS (grey). The axes are colored in red and green for JD and CTD, respectively. Crosslink pairs between JD-CTD, JD-JD and CTD-CTD are shown in a dashed box colored grey, red and green, respectively. Contacts to helix 5 are denoted with H5. **(g**) Histogram of overall R_h_ of DnaJB8 in 1xPBS 285mM NaCl at times 0h (blue), 10h (black), and 20h (red) with arrows indicating R_h_ peaks for each time point. Over time, there is no change in the species of particle sizes <10nm and no appearance of particles ∼100-1000nm. **(h)** Contact map of DnaJB8 crosslinks identified using ADH/DMTMM in the presence of 285mM NaCl. The axes are colored in red and green for JD and CTD, respectively. JD-JD and CTD-CTD crosslinks are shown in dashed boxes colored in red and green, respectively.

We next sought to characterize biochemical properties of DnaJB8-Clover expressed in mammalian cells. DnaJB8-Clover protein was purified using α-GFP nano-bodies^23-24^ (Supplementary Fig. 1c). To gain insight into the topology of DnaJB8, we employed a XL-MS approach to define contacts between different domains ^25-27^. Isolated DnaJB8-Clover was reacted with adipic acid dihydrazide (ADH) and 4-(4,6-dimethoxy-1,3,5-triazin-2-yl)-4-methyl-morpholinium chloride (DMTMM). ADH covalently links carboxylate-carboxylate contacts via a 6-carbon bridge, while DMTMM forms a direct covalent bond between lysine-carboxylate groups through dehydration^26^. Crosslinking treatment of the purified soluble DnaJB8-Clover species revealed predominantly monomers and dimers in the cells, with a trace of larger oligomers (Supplementary Fig. 1d). We identified 21 crosslinks that parsed into 3 regions: JD-JD, CTD-CTD and JD-CTD (Fig. 1d, Supplementary Data 1). The three local JD contacts (Fig. 1d; red box) are consistent with its experimental structure (Supplementary Fig. 1e). Local CTD contacts (Fig. 1d, green box) were also accompanied by inter-domain JD-CTD contacts that localize to helices 2 and 3 of JD (Fig. 1d, grey box). Additionally, we identified a contact between the JD and a putative helix 5 (Fig. 1d, H5) of the G/F domain, as also recently identified in DnaJB6b^17^. Thus, soluble DnaJB8-Clover species isolated from mammalian cells reveal an array of inter-domain interactions, including contacts between the charge complementary JD and CTD.

### DnaJB8 domain contacts are preserved *in vitro*

For a more detailed understanding of DnaJB8 domain architecture *in vitro*, we produced recombinant DnaJB8 as described previously^16^. We first used Dynamic Light Scattering (DLS) to monitor the hydrodynamic radius (R_h_) of DnaJB8 species over time. The scattering data reveals *bona fide* DnaJB8 sizes which begin as a small 4.28±0.82nm species with a small (<%1 by mass) contribution of larger species (>10nm) but over time these small species shift to 5.35±0.22 nm at 10 hours and to 5.77±0.43 nm after 20 hours (Fig. 1e and Supplementary Fig. 1f). Over the time course, a fraction of the soluble small species converted into larger oligomers >10nm (30.6% by mass) with an average R_h_ of 90nm (Fig. 1e and Supplementary Fig. 1f). These findings are consistent with prior studies on DnaJB6b and DnaJB8 showing that they have the capacity to assemble into polydisperse soluble oligomers *in vitro*^16, 18-19, 21, 28^.

Next, we aimed to better understand the topology of DnaJB8 *in vitro* using XL-MS, employing two parallel chemistries; disuccinimidyl suberate (DSS) and ADH/DMTMM on samples after a brief 30 mins of incubation. Consistent with the DLS data at early time points, we observe by SDS-PAGE a ladder of bands indicating the formation of covalent intermolecular contacts dominated by a dimer (Supplementary Fig. 1g). XL-MS analysis of these samples showed only 3 crosslinks in the DSS condition (Fig. 1f, Supplementary Data 1). In contrast, the ADH/DMTMM analysis yielded 24 crosslinks (Fig. 1f, Supplementary Data 1). Importantly, this XL-MS pattern persisted across the DLS time course (Supplementary Fig. 1h and Supplementary Data 1) and closely matches the pairs observed in the assemblies recovered from the mammalian cells including crosslinks from both JD and CTD to H5 (Fig. 1d, H5). The JD crosslinks are consistent with the structure of the domain (Supplementary Fig. 1i)^26^.

The CTD yielded 8 crosslinks (Fig. 1f). Amongst these, the locally-linked regions spanning E208-E211 and K223-K227 are central to the CTD and repeatedly react to peripheral sites. The third cluster of contacts linked the distal JD and CTD through 4 crosslinks (Fig. 1f; JD:CTD). The interdomain CTD crosslink sites are exclusively mediated through acidic amino acids; D212, E209 and E211. Interestingly, the crosslinked amino acids on the JD are all lysines that localize to basic surfaces along helix 1 and 2 (Supplementary Fig. 1j) and overlap the basic Hsp70 binding surface^10^. These XL-MS data identify an intricate network of electrostatic inter-domain interactions in both monomeric and oligomeric DnaJB8.

To further test the apparent role of electrostatically driven interactions, we used a higher ionic strength buffer in an analogous series of experiments. Using DLS we observed a defined species with a 7.75±0.7nm size in 285mM NaCl (Fig. 1g, Supplementary Data 2), which are more expanded compared to species in 150mM NaCl (Fig. 1e, Supplementary Data 2). XL-MS analysis recapitulates the short-range contacts within the JD and CTD domains but the JD:CTD contacts were notably absent (Fig. 1h). To control for reactivity in each condition we compared the frequency of ADH-driven singly reacted modifications, called monolinks. These data show nearly identical numbers of modifications suggesting that the reactivity between these two conditions is nearly identical (Supplementary Fig. 1k). Thus, the disruption of electrostatically driven interactions is accompanied by changes in the domain architecture.

### JD:CTD interaction is mediated by electrostatic contacts

To understand how the JD and CTD domains could be interacting, we used Rosetta modeling guided by XL-MS restraints. We built a starting model by combining the experimental structure of the JD (PDBID: 2DMX) with an *ab initio*-derived model for CTD and the middle domains fully extended. The starting model was then collapsed by applying the JD-CTD crosslinks as restraints (Fig. 2a and Supplementary Fig. 2a-b). The fully expanded monomer collapsed from a predicted R_h_ of 9.27 nm (R_g_, 6.65 nm) to 4.02 nm (R_g_, 2.45 nm) (Fig. 2a). Comparing these values to our DLS radii in 150mM NaCl suggests that the dominant species are likely monomers and dimers. The DLS measurements in 285mM NaCl are consistent with the initial expanded model with the JD:CTD contacts disengaged. Thus, our data support that DnaJB8 exists in solution as small soluble species (4 to 6 nm), dominated by monomer/dimer but with capacity to form larger oligomers over time, both *in vitro* and *in vivo*.

**Figure 2.**
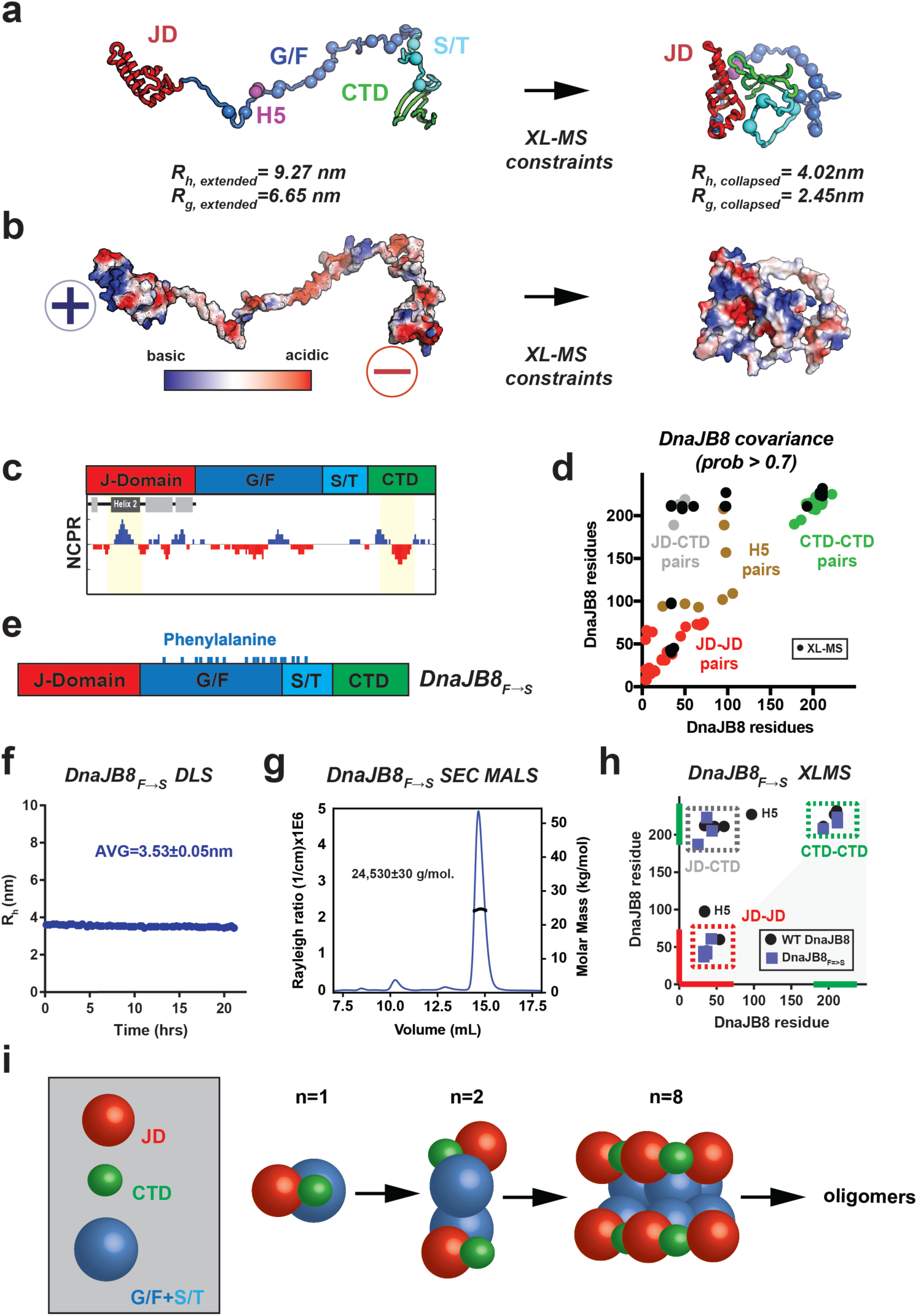
Model for the JD-CTD contacts in a DnaJB8 monomer. (**a**) XL-MS-based refinement of full length expanded DnaJB8 monomer. Cartoon representation of DnaJB8 in fully expanded conformation (left) and collapsed conformation (right), colored by domain as in Fig. 1. Aromatic amino acids in the G/F and S/T domains are shown as spheres and colored according to the domain. Residues in helix 5 (H5) are shown as magenta spheres. Collapsed conformation model was selected from 1,000 Rosetta *ab initio* generated models using a relax protocol. R_g_ and R_h_ values were calculated from the structural model in Rosetta and HYDROPRO, respectively. (**b**) Charge complementary surfaces on the JD and CTD mediate the interaction. Highly acidic potential is shown in red (-sign) and highly basic in blue (+ sign). **(c)** Net charge per residue (NCPR) distribution, defined as the average charge over a 10-residue window, highlights charge complementarity between basic and acidic residues on the JD and CTD, respectively (coloring as in Fig. 1). **(d)** GREMLIN sequence-based covariance analysis identified high confidence covarying amino acids on DnaJB8 that localize within the JD (red), within CTD (green), with H5 (brown) and across JD-CTD (grey). XL-MS links for full length DnaJB8 (black dots) overlap with the covarying regions. Covarying positions localizing to amino acids in G/F domain are shown in brown and co-localize with XL-MS crosslinks. **(e)** Domain map of the DnaJB8_F→S_ mutant, with mutated phenylalanine positions marked by cyan ticks. **(f)** DLS time course of the DnaJB8_F→S_ mutant. The average R_h_ was calculated to be 3.53±0.05nnm. **(g)** SEC-MALS of CTD_170-232_ shows a single peak that was calculated to have a molar mass of 24,530±30 g/mol consistent with a monomer. **(h)** XL-MS contact map showing ADH/DMTMM crosslinks for WT DnaJB8 and DnaJB8_F→S_ mutant. The axes are colored in red and green for JD and CTD, respectively. Crosslink pairs between JD-CTD, JD-JD and CTD-CTD are shown in a dashed box colored grey, red and green, respectively. Contacts to helix 5 in WT DnaJB8 are denoted with H5. **(i)** Schematic of DnaJB8 species observed in solution based on DLS dimensions. Domains are shown as JD (red spheres), CTD (green spheres), and G/F+S/T (light blue spheres). The average R_h_ of DnaJB8_F→S_ (3.53±0.05nnm) and DnaJB8 *ab initio* Rosetta model (4.02nm) are assigned to the monomer. The R_h_ of WT DnaJB8 begins as a 4.28nm species and grows to 5.77nm over 20hrs. Size and volume estimates from the structural models suggest DnaJB8 exists as small species ranging from a monomer to octamer likely dominated by a dimer and over time maturing into large oligomers.

Guided by the constraints, the final model “docks” the JD onto the CTD placing a putative acidic surface on the CTD in contact with the basic surface on the JD (Fig. 2a and Supplementary Fig. 2b) and additionally bringing H5 in proximity to both the JD and CTD as similarly observed for DnaJB6b (Supplementary Fig. 2c) ^17^. The CTD has proximal basic surfaces that flank its acidic surface, generating a characteristic alternating charge pattern that is inverted on the JD (Fig. 2c). Mapping sequence conservation onto the Rosetta-generated model, we find that these JD:CTD contacts are largely conserved (Supplementary Fig. 2d). In a co-evolution analysis using the Gremlin algorithm^29-31^ we identified amino acid positions that covary. Not only did we observe many amino acid pairs that covary between the JD and CTD as well as H5, but our XL-MS pairs overlap with these covarying positions (Fig. 2d). The similarity between the predicted covarying contacts, conservation, and the XL-MS experimental contacts strengthens our DnaJB8 JD-CTD model, and suggests that XL-MS can detect functionally important interaction sites.

### JD:CTD contacts are present in monomeric DnaJB8

In our “collapsed” monomer structural model the 17 phenylalanine residues in the G/F and S/T domains were predicted to be in part solvent exposed (Fig. 2a, spheres). We hypothesized that these aromatic residues may play a role in DnaJB8 assembly and engineered a mutant, in which all G/F- and S/T-region phenylalanine residues were mutated to serine residues (Fig. 2e, herein DnaJB8_F→S_). Using our DLS and XL-MS pipeline, we evaluated assembly of DnaJB8_F→S_. By DLS, the DnaJB8_F→S_ mutant remained stable as a 3.53±0.05 nm species over 21 hours (Fig. 2f, Supplementary Data 2). SDS-PAGE of crosslinked DnaJB8_F→S_ also showed no intermolecular crosslinks. (Supplementary Fig. 2e). Size Exclusion Chromatography Multi-Angle Light Scattering (SEC-MALS) analysis on DnaJB8_F→S_ revealed it to be a monomer with a molecular weight of 24,530±30 g/mol (Fig. 2g). These data support that phenylalanine residues in the G/F and S/T domains play a role in higher order assembly. Next, we used XL-MS to test whether this DnaJB8_F→S_ monomer maintained the intramolecular JD and CTD contacts observed in WT DnaJB8 (Fig. 2h). Analysis of the crosslinked DnaJB8_F→S_, revealed identical local crosslinks within JD and CTD and also detected 3 crosslinks between the JD and CTD. Interestingly, in DnaJB8_F→S,_ the H5 crosslinks to JD were absent, consistent with requirement of a phenylalanine in H5 for binding to the JD (Fig. 2h, H5). The presence of the JD-CTD crosslinks in a monomeric mutant marks them to represent *intra*molecular JD:CTD interactions.

We can now use the experimental DLS radii with our structural models to more accurately infer the dimensions of the small soluble DnaJB8 species (Fig. 2i). At the start of the WT DnaJB8 DLS time course, we observed an initial population of polydisperse particles with an average radius of 4.28±0.82nm (Fig. 1e). The DnaJB8_F→S_ mutant showed a radius of 3.53±0.05nm with a very narrow monodisperse distribution, further supporting our model of a monomer “collapsed” by JD:CTD interactions. Based on a model proposed by Marsh and Forman-Kay^32^, we also estimate that a monomeric 232-residue DnaJB8 protein should have a size of 3.98 nm. These data support our analysis that WT DnaJB8 at first adopts primarily a monomer/dimer distribution that has the capacity to then assemble into large oligomers. In contrast, the larger DLS R_h_ values measured for DnaJB8 in 285mM NaCl (Fig. 1g), are a result of the loss of the JD:CTD contacts yielding a small oligomer mediated by aromatic contacts.

### ssNMR on DnaJB8 oligomers reveals regions of disorder and order

For additional insight into their molecular structure and dynamics, magic-angle-spinning (MAS) ssNMR was performed on the hydrated oligomers of U-^13^C,^15^N-labeled DnaJB8. MAS ssNMR of hydrated protein assemblies allows for the site- and domain-specific detection of mobility and (secondary) structure, even in presence of disorder and heterogeneity. 1D and 2D ssNMR spectra of the DnaJB8 oligomers feature many broad peaks, with linewidths up to 0.38 kHz, consistent with an oligomeric assembly displaying structural disorder (Fig. 3a; Supplementary Fig. 3a-d). However, strikingly, distinct subsets of narrow peaks are also detected, with linewidths of 0.1 to 0.2 kHz (Fig. 3c-d left). These ssNMR experiments employ the cross-polarization (CP) technique, in which observable residues must be rigid or immobilized^33^. In INEPT-based ssNMR, which is selective for highly dynamic segments, the oligomers show little signal^33-36^ (more below). Then, the observed narrow signals in CP spectra must originate from an immobilized, well-ordered subset of DnaJB8 residues. These narrow signals are from amino acid types^37^ in the JD while the broad peaks are dominated by signals from residues common in other domains (Supplementary Table 1). The former also reflect mostly α-helical structure, while the latter are mostly random coil and β-sheet ^38^. With known chemical shifts of the DnaJB8 JD in solution, we prepared a synthetic 2D spectrum (Supplementary Fig. 3b red) that has a striking correspondence to the narrow ssNMR peaks (Supplementary Fig. 3b black), such that we tentatively assign those to residues in helix 2 and helix 3. The 2D ^15^N-^13^Cα (NCA) ssNMR spectrum showed a similar alignment between narrow peaks and JD signals in solution (Supplementary Fig. 3d). These CP-based 2D spectra also feature strong peaks from immobilized charged side chains (Lys, Arg, Asp, Glu; Supplementary Figure 3e-f), which is consistent with their involvement in salt-bridge interactions predicted by the XL-MS analysis above.

**Figure 3.**
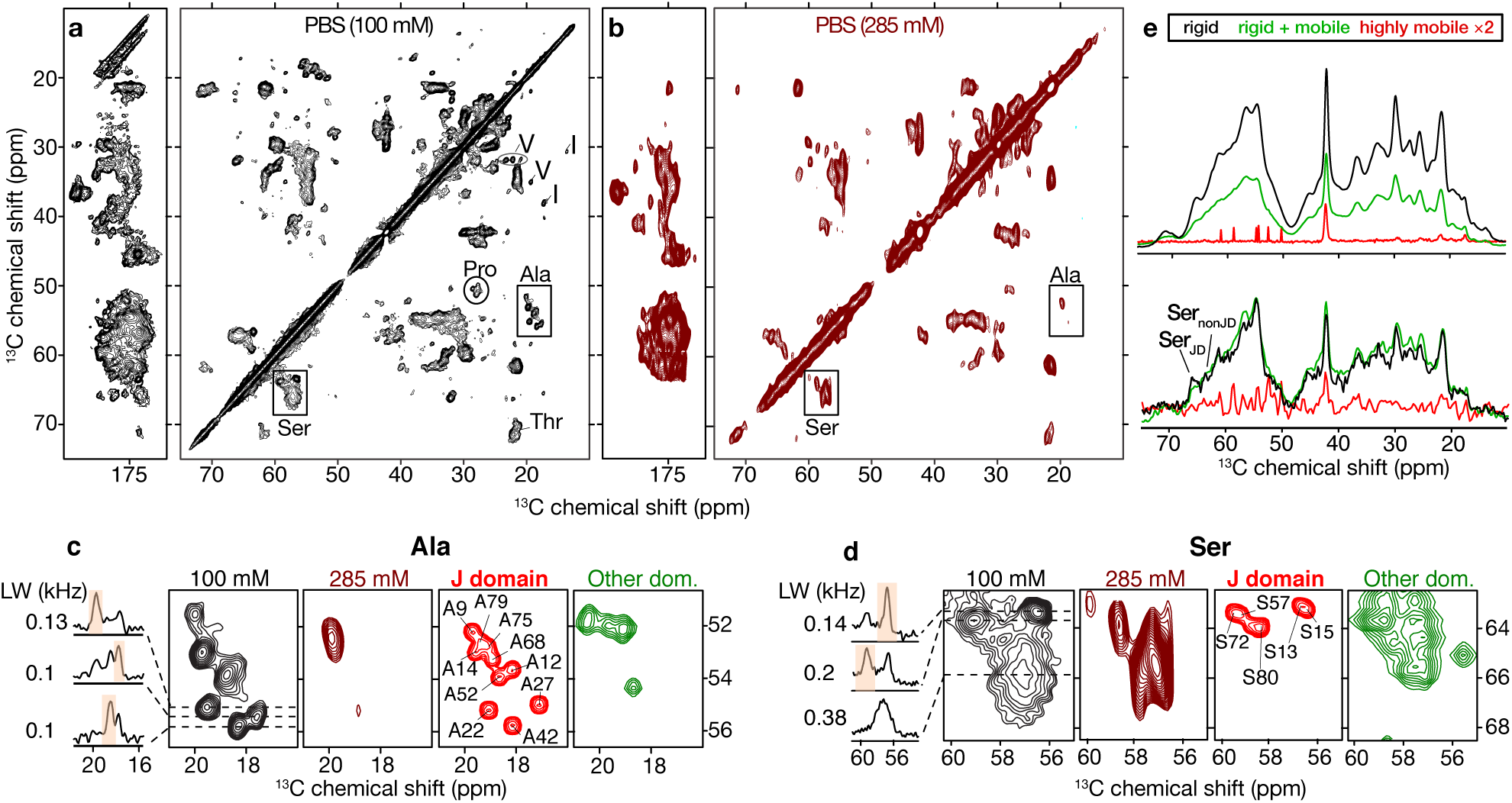
Solid-state NMR of DnaJB8 oligomers at physiological and high ionic strength. **(a)** 2D ^13^C-^13^C ssNMR spectrum of U-^13^C,^15^N-labeled DnaJB8 oligomers in PBS (100 mM NaCl), using 25 ms DARR mixing. **(b)** Corresponding 2D ssNMR spectrum in PBS with 285mM NaCl. **(c-d)** Boxed Ala and Ser regions from panels “a” and “b”. In PBS, the experimental Ala and Ser peak patterns (black) are well resolved and similar to those expected for folded JD in solution (red). At elevated ionic strength (brown) these narrow peaks are missing. Green spectra (right) represent simulated signals predicted for our models of the non-JD domains, shown with enhanced broadening reflecting the heterogeneity seen in the experiments. 1D spectra on far left show slices through the experimental 2D data, with selected peak widths (in kHz). (**e**) ^13^C 1D spectra, in PBS (top) and with 285 mM NaCl (bottom), that show rigid residues (black, CP), rigid and mobile residues (green, SPE) and only mobile residues (red, INEPT). See also text and Supplementary Fig. 3.

In absence of experimental solution NMR data for other domains, we predicted estimated spectra based on our structural models (Fig. 3c-d green; Supplementary Fig. 3c)^39^. These peak patterns qualitatively resemble the broad signals in our 2D ssNMR data. A particular strength of MAS ssNMR of hydrated proteins is the ability to gauge local and global dynamics. Single pulse excitation (SPE) and refocused INEPT spectra, which enhance the more dynamic parts of samples^35-36^ show surprisingly little evidence of flexible residues (Fig. 3e top red). Indeed, the main INEPT signal (∼42 ppm) is just from solvent-exposed Lys side chains and lacks evidence of flexible protein regions (even from the S/T or G/F regions). Given that the 1D CP and SPE spectra (Fig. 3e top) look similar, with higher signal intensities in the former, the different domains of the protein actually must have a similar degree of mobility and all be mostly immobilized, without flexible regions. Combined, the ssNMR data reveal oligomers that are heterogeneous in structure but lack extended flexible domains. In other words, the central G/F and ST domains are heterogenous, but also immobilized within the oligomers, consistent with the abovementioned role of their Phe residues in driving oligomer assembly. Uniquely ordered are parts of the JD (residues in helices 2 and 3; Supplementary Fig. 3g,h), which show up as well-folded and immobilized.

### Interaction sites from ssNMR

MAS ssNMR studies of DnaJB8 oligomers in PBS buffer with 285mM NaCl (analogous to the studies above) are shown in Fig. 3b. The 2D spectrum reproduces the broad signals of the immobilized oligomers, but the narrow JD peaks are now strikingly absent. Comparing CP and SPE ssNMR spectra (Fig. 3e bottom), there is an increase in overall mobility. Notably, no new “flexible” ssNMR signals were identified by INEPT ssNMR. We attribute the loss of JD signals in CP-based spectra to increased mobility due to disruption of long-range electrostatic interactions, while the lack of INEPT peaks tells us the JD is still folded and partly immobilized by covalent attachment to the overall assembly. In other words, the JD is invisible due to intermediate timescale dynamics^33, 40^. Since the broad signals from the other domains are preserved, it appears that the core architecture of the oligomers persists, consistent with aromatic and hydrophobic interactions.

### Isolated JD and CTD are folded and monomeric

To further characterize the JD and CTD interaction, we produced isolated JD (herein JD_1-77_) and CTD (herein CTD_170-232_) (Fig. 4a). SEC analysis of JD_1-77_ and CTD_170-232_ revealed monodispersed peaks (Fig. 4b). SEC-MALS determined each domain to be monomeric with a molecular weight of 10,220±220 g/mol and 8,376±14 g/mol for JD_1-77_ and CTD_170-232_, respectively (Supplementary Fig. 4a-b). Also, by DLS we measured the JD_1-77_ R_h_ to be 2.31±0.13 nm and the CTD_170-232_ to be 1.71±0.02 nm, with both stable over 15 hours (Supplementary Fig. 4c, Supplementary Data 2). We again employed XL-MS to probe the individual domains and compare them to full-length protein. On an SDS-PAGE gel, the crosslinked JD_1-77_ and CTD_170-232_ remained monomeric following crosslinking (Fig. 4c). XL-MS analysis yielded 4 crosslinks for JD_1-77_ and 6 crosslinks for CTD_170-232_ (Fig. 4d, Supplementary Data 1). The identified crosslinks revealed good agreement between the local domain crosslinks observed in the full-length DnaJB8 and the isolated domains (Fig. 4d, Supplementary Fig. 4d).

**Figure 4.**
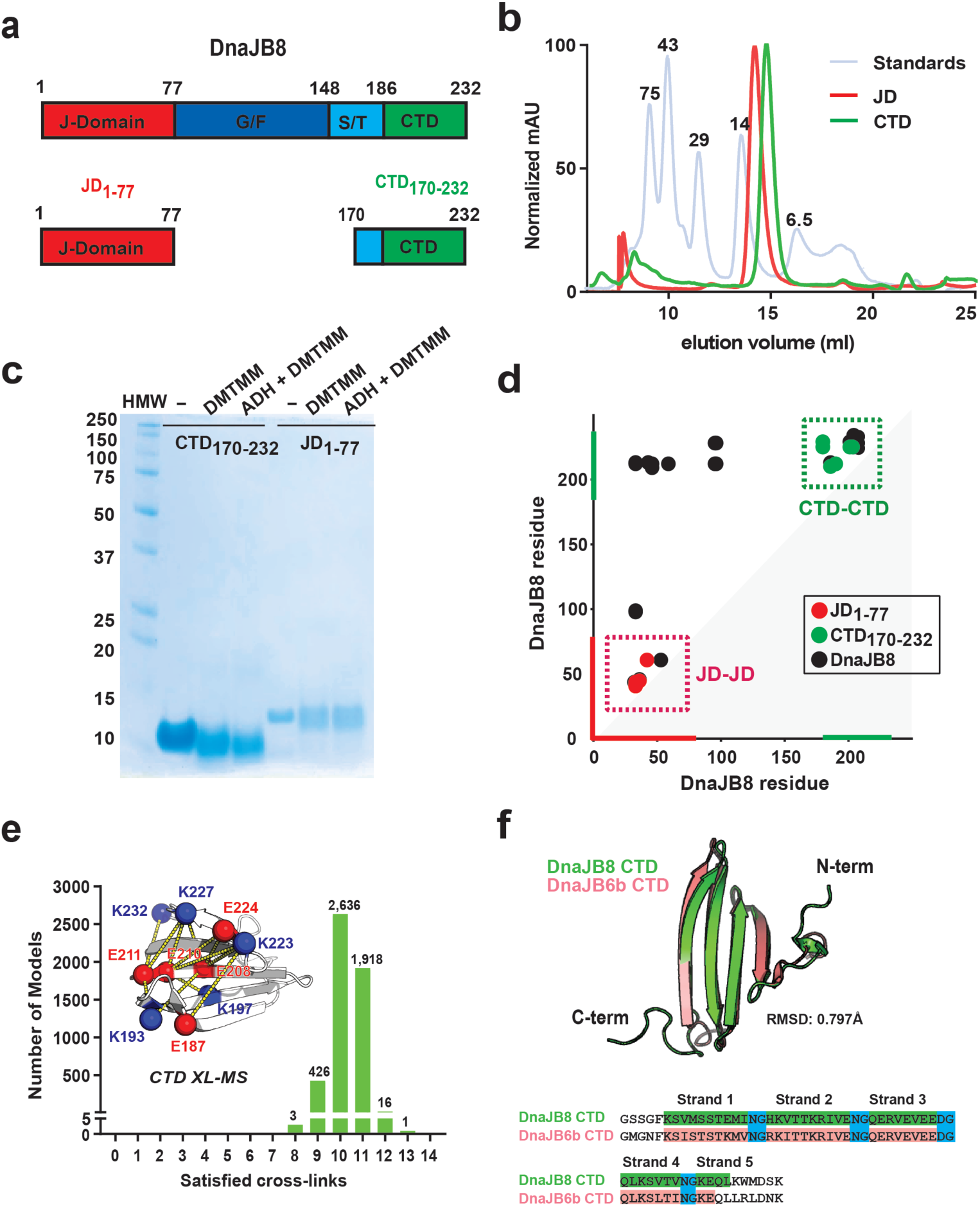
Isolated JD and CTD proteins are monomeric. **(a)** Cartoon schematic for the full length DnaJB8 and domain fragments JD_1-77_ and CTD_170-232_. **(b)** Representative SEC profiles of JD_1-77_(red), CTD_170-232_(green) and LMW standards (blue). JD_1-77_ and CTD_170-232_ elute at apparent molecular weights of 14kDa and 6.5kDa, respectively. **(c)** SDS-PAGE coomassie gel of crosslinked JD_1-77_ and CTD_170-232_ reacted with either DMTMM only or DMTMM with ADH. **(d)** Contact map of ADH/DMTMM crosslinks identified for JD_1-77_(red), CTD_170-232_(green) and full length DnaJB8 (black). The axes are colored in red and green for JD and CTD, respectively. Crosslink pairs between JD-JD and CTD-CTD are shown in dashed boxes colored in red and green, respectively. **(e)** Histogram of the number of intra-domain crosslinks that are consistent with crosslink chemistry geometry (“satisfied”) in the ensemble of 5000 models. One model satisfies 13 out of 14 possible crosslinks identified in our experiments. Crosslinks are mapped onto best matching CTD structural model (inset), shown in white cartoon representation. Sites of crosslink are shown as red or blue spheres, for D/E and K, respectively. Dashed yellow lines connect linked amino acid pairs. **(f)** Overlay of our DnaJB8 CTD model generated by *ab initio* ROSETTA (green) with the published DnaJB6bΔST CTD (salmon) (PDB:6U3R). The CTD sequences of DNAJB8 and DNAJB6 are shown with each β-strand highlighted and conserved NG and DG turns in blue.

We built an ensemble of models for the CTD_170-232_ using *ab initio* ROSETTA^40^. The calculated R_h_ for the structural ensemble was consistent with the DLS measurement of 1.7 nm (Supplementary Fig. 4g). The models formed a low contact order 5-stranded β-sheet topology and the R_h_ variation can be attributed to the more flexible termini (Supplementary Fig. 4g, inset). Circular dichroism (CD) on the CTD sample yields spectra consistent with a predominantly β-sheet content, as predicted by our model (Supplementary Fig. 4e-f). We mapped the 14 CTD-derived crosslinks from across experiments onto the monomeric ensemble, finding that a majority of structures explain 10-11 crosslinks but only a single model explains 13 of 14 (Fig. 4e). These crosslink pairs map onto each face of the β-sheet and the distances are compatible with the geometry of the crosslinking chemistry. The crosslinks that fall outside of the distance cutoff localize to the more dynamic C-terminus of CTD (Fig. 4e and Supplementary Fig. 4g, inset) at positions K227 and K223. The CTD topology is defined by four β-turns stabilized by conserved asparagine/aspartate-glycine sequences (N/DG) and overlays well with the DnaJB6b CTD (Fig. 4f)^17^. Thus, our data support that both the JD_1-77_ and CTD_170-232_ domains are folded, monomeric, and do not have intrinsic assembly properties.

### Basic surface on JD drives interaction with CTD

In our experiments on the full-length DnaJB8 oligomers we observed that the JD and CTD interact through complementary electrostatic surfaces. We further probed this interaction by mixing the individual JD_1-77_ and CTD_170-232_ domains *in vitro* (Fig. 5a). We incubated Flourescein(FITC)-labelled JD_1-77_ with a series of CTD_170-232_ concentrations and measured binding affinity using a fluorescence polarization (FP) assay. The resulting binding curve revealed that the JD_1-77_ binds to the CTD_170-232_ with 0.542±0.071 μM affinity (Fig. 5a; bottom). Parallel experiments using Microscale Thermophoresis revealed a K_d_ of 5.36±1.15 μM (Supplementary Fig. 5a) suggesting that the affinity is in the low micromolar range. JD_1-77_ and CTD_170-232_ domains were mixed together to form the complex and analyzed using XL-MS. We identified 6 local crosslink pairs consistent with pairs observed in full-length DnaJB8 and the isolated JD_1-77_ and CTD_170-232_ samples (Fig. 5b). Importantly, we also reconstitute 4 intermolecular contacts between the JD_1-77_ and CTD_170-232_ observed in full length DnaJB8 experiments. However, an increased variance in the crosslink profile may indicate that the missing proximal sequences help define the proper architecture of the full-length DnaJB8 oligomers. Solution NMR-based chemical shift perturbation (CSP) mapping was used to identify the JD_1-77_ surface that interacts with the CTD_170-232_ (Fig. 5c, Supplementary Fig. 5b-c). Titration of increasing amounts of unlabeled CTD_170-232_ into ^15^N-labeled JD_1-77_ produced fast exchanging concentration-dependent CSPs in a specific subset of peaks (Fig. 5d-e); 33 peaks were strongly perturbed (>0.2 p.p.m.) and another 40 peaks were perturbed weakly (>0.1 p.p.m.). Among the strongly perturbed peaks, 27 residues have solvent accessible side chains of which fifteen are charged, with a notable overlap with the basic face in helix 2 that is implicated in Hsp70 binding (Fig. 5f-h). Other residues that show strong perturbations are basic and acidic residues that wrap around the exterior of the JD loop and helix 3, and residues on the charged face of helix 4. While a few other hydrophobic residues also show strong perturbations, all are in close proximity to charged residues along each helix. Given the small size of the JD_1-77_, it is likely that residues in the core behind the basic surface involved in the interaction experience changes in chemical shift.

**Figure 5.**
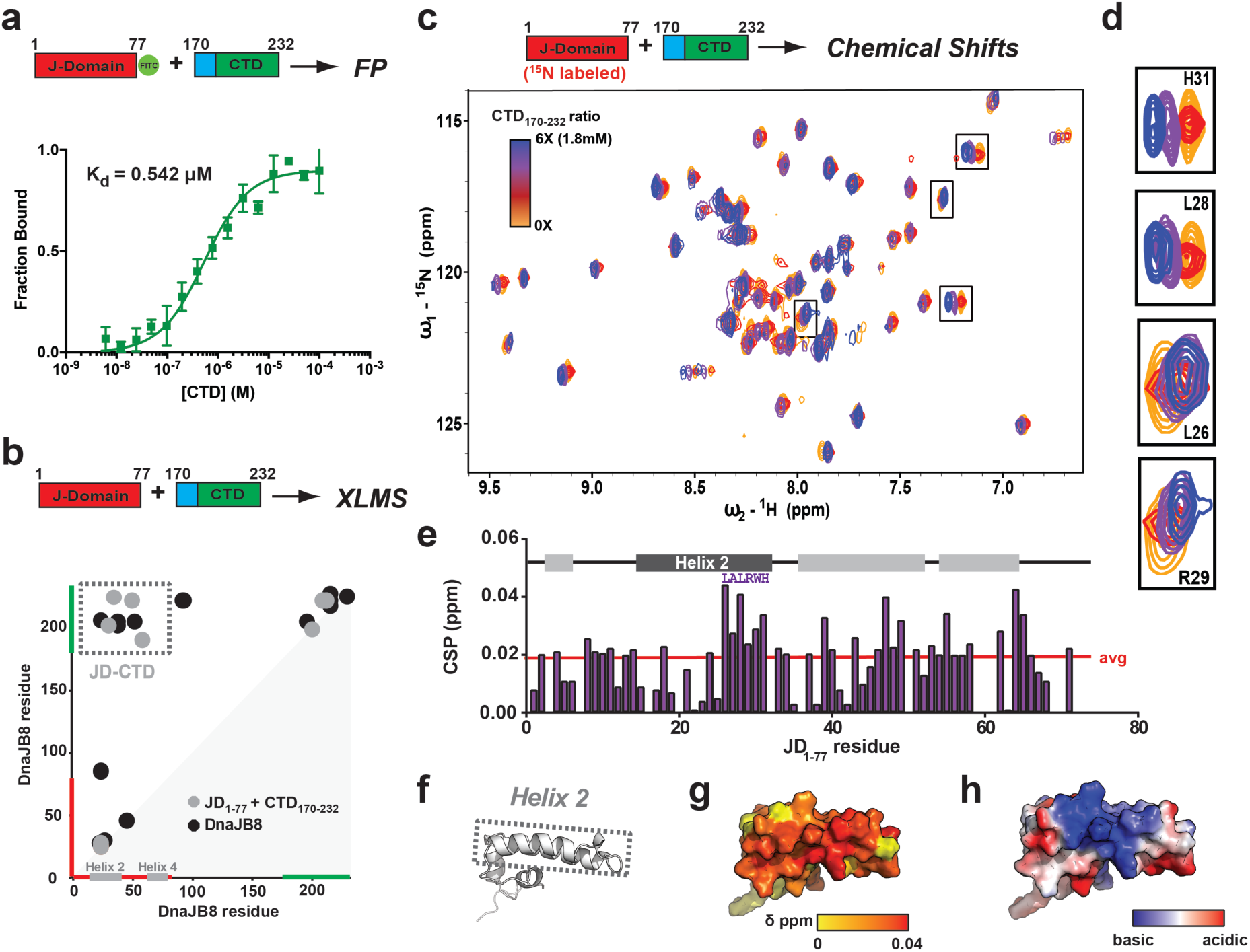
JD and CTD interact through charge complementary surfaces. **(a)** Schematic of the JD_1-77_-FITC (FITC dye is shown as a green circle) and CTD_170-232_ constructs used in fluorescence polarization (FP) experiments. FP titration measuring interaction between JD_1-77_-FITC and a concentration range of unlabeled CTD_170-232_. FP experiments were performed in triplicate and shown as averages with standard deviation. **(b)** Schematic of the JD_1-77_ and CTD_170-232_ constructs used in the XL-MS experiments. Contact map of ADH/DMTMM crosslinks identified from an incubated JD_1-77_ and CTD_170-232_ sample (grey) and full length DnaJB8 (black). The axes are colored in red and green for JD and CTD, respectively. Helix 2 and 3 are shown in grey on the x-axis. Crosslink pairs between JD-CTD are shown in a dashed box colored in grey. **(c)** Schematic for the solution NMR chemical shift experiment with U-^15^N JD titrated with unlabeled CTD. HSQC solution NMR spectrum of 300μM ^15^N-labelled JD_1-77_ against a titration of CTD_170-232_: 0x (yellow), 1x (red), 3x (purple), 6x (blue). DnaJB8 JD peak assignments were transferred from deposited data (BMRB:11417). **(d)** Insets of peaks with highest observed chemical shifts: H31, L28, L26, R29. Coloring as in **c. (e)** Histogram of chemical shift perturbations (CSP) from 3x CTD experiment by residue. Average CSP of ∼0.019ppm is denoted by the red line (excludes prolines). The sequence of last 6 residues of helix 2 are marked above their respective peaks. **(f)** DnaJB8 JD structure illustrating the location of helix 2 (pdbid:2DMX). **(g)** Mapping CSP values onto the DnaJB8 JD structure, shown in surface representation and colored according to Δδ from low (0.0 ppm) in yellow to high (red; 0.04 ppm). **(h)** Electrostatic potential mapped onto DnaJB8 JD structure shown in surface representation. Highly acidic potential is shown in red and highly basic in blue.

### JD-CTD interaction competes with Hsp70 binding

The recent X-ray structure of the DnaK:DnaJ complex revealed a conserved charge-based interaction between the basic surfaces on the JD of DnaJ and an acidic surface on DnaK^10^. Using this complex as a template, we modeled the binding interface of the human Hsp70 (HspA1A)^41^ and the JD of DnaJB8^42^ (Fig. 6a-c). The basic surface on the DnaJB8 JD (Fig. 6b) contacts the conserved acidic surface on HspA1A (Fig. 6c, Supplementary Fig. 6a-b). Thus, conserved electrostatic contacts are likely to play a key role in the interaction between Hsp70 and Hsp40.

**Figure 6.**
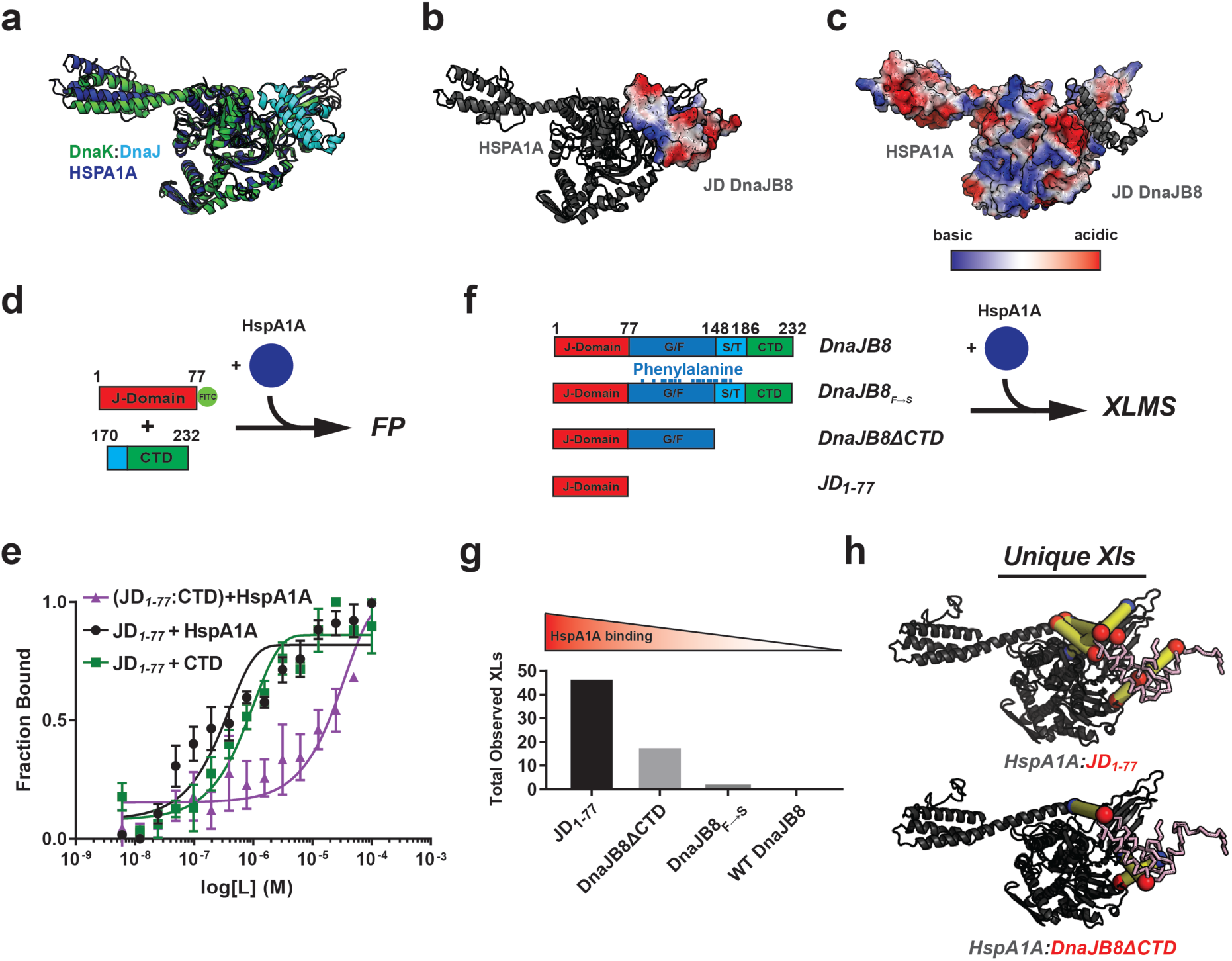
CTD and HSPA1A compete for the same basic binding surface on DnaJB8aD. **(a)** Structural superposition of a representative HspA1A structural homology model (blue) with a crystal structure of DnaK-DnaJ (green and cyan, respectively; PDBID: 5NRO) shows good agreement. **(b-c)** Electrostatic surface potential of DnaJB8 JD docked into the JD binding site on HspA1A (shown in black cartoon representation). Basic surface on helix 2 docks onto the HspA1A surface. Electrostatic surface potential of HspA1A with docked DnaJB8 JD in black cartoon representation. The HspA1A surface presents an acidic face that complements the basic DnaJB8 JD surface. Highly acidic potential is shown in red and highly basic is shown in blue. **(d)** Experimental workflow used to determine competition between Hsp70 and CTD_170-232_ for JD_1-77_-FITC binding (dye shown as a green circle). **(e)** Normalized FP binding curves measuring affinity between fluorescent JD and added CTD (green) or added Hsp70 (grey). Preincubation with CTD followed by addition of Hsp70 (purple) shows delay in binding consistent with a competitive binding model. FP experiments were performed in triplicate and shown as averages with standard deviation. **(f)** XL-MS-based experimental workflow used to determine contribution of CTD to regulate JD binding to Hsp70. WT DnaJB8, DnaJB8ΔCTD, DnaJB8_F→S_ and JD_1-77_ DnaJB8 variants were used to form complexes with HspA1A. (**g**) Summary of total intermolecular crosslinks identified across three XL-MS experiments between the JD and HspA1A for four complexes: JD_1-77_:HspA1A, DnaJB8ΔCTD:HspA1A, WT DnaJB8:HspA1A and DnaJB8_F→S_:HspA1A. (**h**) Unique intermolecular crosslinks identified across three datasets in the JD_1-77_:HspA1A and DnaJB8ΔCTD:HspA1A complexes mapped onto the JD-HspA1A model. JD is shown in pink ribbon representation and HspA1A in black cartoon representation. Sites of crosslink are shown as red or blue spheres for aspartic/glutamic acid and lysine, respectively. Yellow lines connect linked amino acid pairs.

The conserved HspA1A-JD electrostatic contacts (Fig. 6b-c) that overlap with the JD-CTD contact sites lead us to hypothesize that the observed JD-CTD interactions could interfere with Hsp70 binding. To test this hypothesis, we employed a competition experiment leveraging our FP binding assay to discriminate the JD-CTD and JD-HspA1A complexes (Fig. 6d). We determine a 0.257±0.029μM affinity for the JD:HspA1A interaction, consistent with values in the literature^43^ and similar to the JD:CTD interaction (Fig. 6e, black and green, respectively). Due to the size difference between HspA1A (70 kDa) and the CTD (8.7 kDa) their respective complexes with tagged JD plateau at different polarization values (Supplementary Fig. 6c, black and green, respectively). Leveraging this difference, we designed a binding experiment to measure the competition of HspA1A and CTD binding to the JD. FITC-labeled JD was preincubated with 3 μM CTD, followed by a titration with HspA1A. The pre-titration FP signal was consistent with formation of the JD-CTD complex, which persisted until HspA1A concentrations of 3.125uM when the signal shifted to that of a JD-HspA1A complex (Supplementary Fig. 6c, purple). Normalization of the data reveals a ∼25-fold decrease in the apparent binding constant between JD:HspA1A when preincubated with CTD (Fig. 6e). To further test the inhibitory role of CTD on recruitment of Hsp70, we used XL-MS to measure the frequency of HspA1A and JD contacts across a set of complexes formed between HspA1A and WT DnaJB8, JD_1-77_, DnaJB8_F→S_ and DnaJB8ΔCTD missing the CTD (Fig. 6f). Across three experiments, we detected no crosslinks between the Hsp70 and the JD in WT DnaJB8 and only 2 in DnaJB8 _F→S_ (Fig. 6g, Supplementary Data 1). In contrast, in the HspA1A:JD_1-77_ and HspA1A:DnaJB8ΔCTD complexes, we identified 47 and 14 total crosslinks between the JD and HspA1A, respectively (Fig. 6g). All identified pairs are consistent with the structural model (Fig. 6h and Supplementary Fig. 6d). These data support that the robust JD:CTD engagement seen in WT DnaJB8 and even the monomeric DnaJB8_F→S_ (Supplementary Fig. 6e) prevents HspA1A interaction with the JD domain and deletion of the CTD releases the inhibitory effect (Fig. 6g). Thus, the DnaJB8 JD uses a basic surface to bind an internally encoded CTD via an acidic surface that directly inhibits HspA1A binding.

## DISCUSSION

### Modeling the shape of DnaJB8

DnaJB8, like DnaJB6b, has the capacity to assemble into soluble oligomers. We used a combination of protein engineering, solution scattering data and modeling to understand the shapes of DnaJB8 in solution. Using our XL-MS data, we collapsed a DnaJB8 structural model around the JD:CTD interaction and thus obtained a structural model that fit the average R_h_ of the monomer measured by DLS. Based on the fold of this monomeric model, we hypothesized that aromatic amino acids in the central G/F and S/T domains would be exposed and thus could mediate self-assembly into oligomers. Indeed, mutagenesis of aromatic residues yielded a stable monomeric variant of DnaJB8 in agreement with our collapsed structural model with engaged JD:CTD contacts. This is further supported by good agreement between the R_h_ of our collapsed structural model, DLS data, and values derived from the Marsh and Forman-Kay model^32^. An intriguing question relates to whether our models may also be applicable to DnaJB6b. At this time a direct comparison is difficult given the known structural and functional differences of the proteins and the lack of analogous experimental data, especially on the larger oligomers of DnaJB6b. Our collective data highlights the power of our multipronged approach to derive the base unit of a DnaJB8 monomer, which employs exposure of aromatic residues to mediate assembly through nonpolar surfaces into larger oligomers.

### Functional role of the CTD in DnaJB8

We combined XL-MS and NMR in the solid and solution states to probe DnaJB8 inter-domain interactions. One of the most striking features was a newly identified interaction between the distal JD and CTD driven by electrostatics. This interaction was perturbed by the addition of salt, but maintained following mutagenesis of aromatic amino acids in the central domains. Since analysis of the isolated JD and CTD showed a reduced mutual association, there nonetheless is a distinct role for the intervening domains in the JD-CTD interaction. Our combined data show that the DnaJB8 S/T and G/F domains are not behaving as “flexible linkers”^17^ and that their aromatic residues are central in the homo-oligomerization process. On their own, both JD and CTD are surprisingly resistant to self-assembly. These findings are distinct from published reports on DnaJB6b, where the CTD appears to drive oligomerization, which may relate to sequence divergence in the six C-terminal CTD residues between DnaJB8 and DnaJB6b^17-18^. Nonetheless, our modeled CTD structure, featuring a pleated β-sheet topology absent of a hydrophobic core, is identical to its recently reported DnaJB6b counterpart^17^. Interestingly, outside of inter-strand hydrogen bonding and polar side-chain contacts, it is not clear what forces stabilize this domain. This may explain the CTD heterogeneity (unlike the JD) seen by ssNMR. The CTD topology resembles the charged β-sheet surface on Hsp70 that is known to interact with the JD^10^. While our reconstitution of the JD:CTD interaction using isolated domains indicates that the CTD alone can bind the JD, we cannot exclude that helix 5 can contribute to this interaction to regulate Hsp70 function. It is worth noting that lysine residues in the DnaJB8 and DnaJB6b CTD can be acetylated and deacetylated (via Histone Deacetylases (HDACs)) to modify these proteins’ self-assembly and function, which may involve changes in the Lys-mediated JD interactions^16, 20^. The CTD architecture is conserved in a broader subset of B family member Hsp40s^18^. We speculate that the CTD in these DnaJB family members similarly serves a regulatory role in which post-translational modifications could alter the affinity for the JD, and thus indirectly alters oligomerization or Hsp70 recruitment.

### Implications for Hsp70 recruitment and substrate binding

Aside from suppressing protein aggregation on its own^16, 18, 20^, DnaJB8 also recruits Hsp70 for processing of bound substrates. Our current findings hint at an intriguing possibility that auto-inhibitory interactions of the Hsp70-binding JDs within the DnaJB8 oligomer could be involved in substrate-binding-coupled Hsp70 recruitment. In the non-stressed native state, DnaJB8 forms soluble oligomers in which the JD is engaged in electrostatic interactions and thus not available for Hsp70 binding as supported by our experiments (Fig. 6g; Fig. 7a). We hypothesize that substrate binding could allosterically disrupt the JD-CTD interaction, exposing the Hsp70-binding HPD motif of the JD (Fig. 7b). This would enable the recruitment of Hsp70 to the loaded DnaJB8 protein. Aromatic-driven oligomeric assembly of DnaJB8 may be related to the formation of liquid-liquid phase separated assemblies in other proteins containing similar arrangements of phenylalanine residues^44^. We propose that the more hydrophobic elements of the G/F and S/T domains form the oligomer core, with the CTD and JD remaining relatively surface exposed. Thus, it may be possible to recruit Hsp70 to different DnaJB8 species. Our data on the DnaJB8_F→S_ mutant illustrate that the JD:CTD interaction exists in the monomeric base unit suggesting that this interaction is present across the polydisperse distribution of DnaJB8 species. Although we as yet lack detailed information supporting a substrate-triggered modulation of the JD:CTD interaction, our results offer some hints toward a possible molecular mechanism for such a coupling. In *in vivo* and *in vitro* XL-MS experiments, negatively charged residues in helix 5 in the G/F domain interact with both the JD and CTD (Fig. 1). We also saw a change in JD:CTD affinity in the absence of the central domains (Fig. 5). Finally, other studies on DnaJB6b have identified the S/T domains as substrate binding domains^16, 20-21^. Future mechanistic and structural studies on DnaJB8 and other complex chaperones including DnaJB6b and their interactions with substrates will reveal the interplay between oligomer dynamics, post-translational modifications, substrate binding, and recruitment of Hsp70.

**Figure 7.**
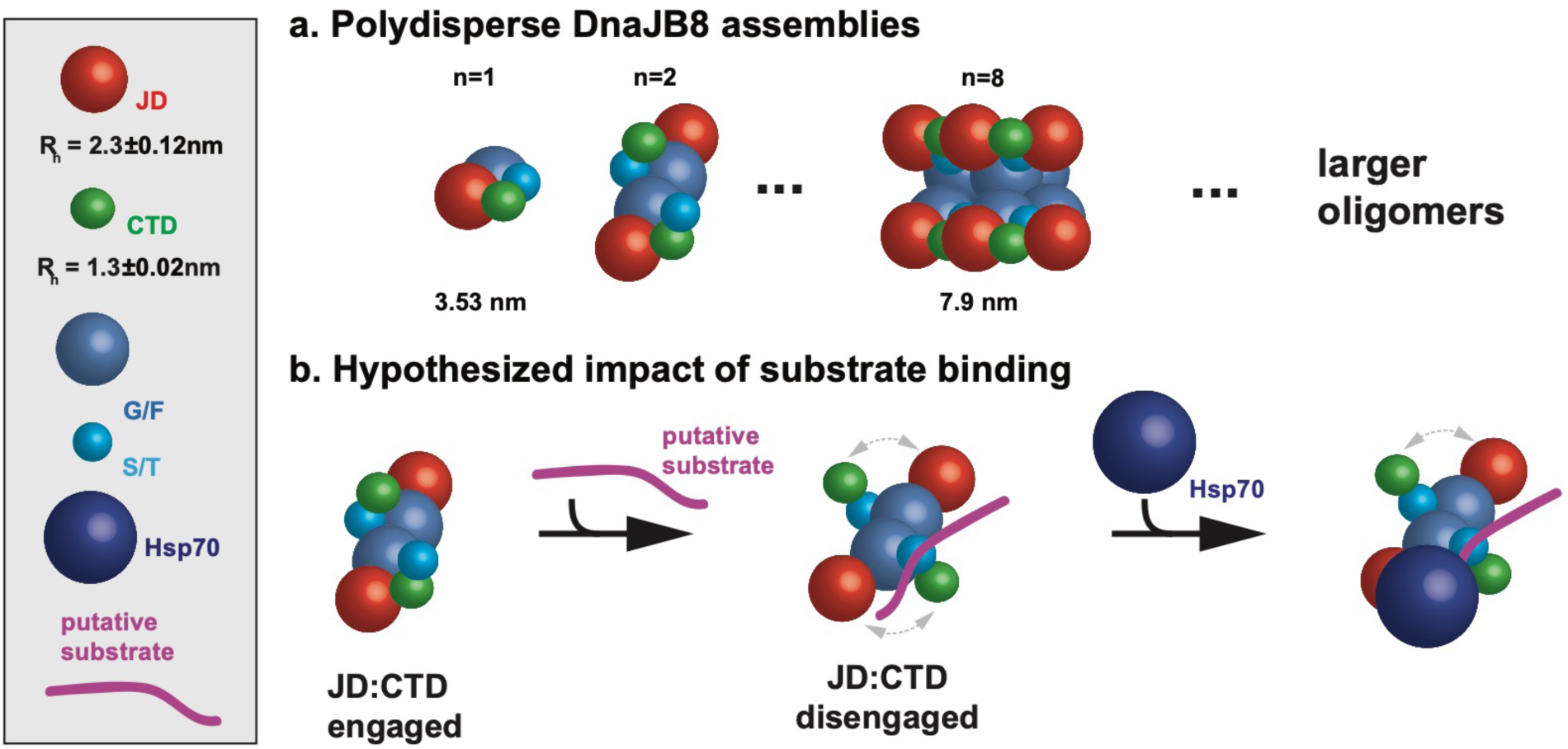
Proposed model for DnaJB8:HspA1A:Substrate relationship. Schematic of proposed DnaJB8 model. Domains are shown as JD (red spheres), CTD (green spheres), G/F (blue spheres), S/T (light blue spheres), and also HspA1A (dark blue spheres) and substrate (purple line) are shown. DnaJB8 domain sizes are displayed scaled to the relative R_h_ values derived from DLS experiments (HspA1A not drawn to scale). **(a)** DnaJB8 forms a fundamental oligomeric species through aromatic contacts in the G/F and S/T domains ranging from monomer to octamer. **(b)** The JD:CTD engaged state, where the JD is stabilized by CTD and helix 5 (G/F) contacts, can form larger polydisperse oligomers (>100nm). The JD:CTD disengaged state (bottom) is needed to engage with HspA1A. We illustrate our hypothesis where substrate binding may allosterically disrupt the JD:CTD interaction to allow the recruitment of HspA1A to the freed JD-CTD binding face, enabling subsequent handoff of substrate to HspA1A.

## Materials and Methods

### Sequence and structural analysis of DnaJB8, DnaJB6b and HspA1A

Analysis of protein sequences (including the net charge per residue; NCPR) was performed using Local CIDER^45^. An ensemble of 1000 HspA1A homology models was produced using *ab initio* Rosetta using the DnaK (PDB:5NRO) conformation as a template^10^. Briefly, the HspA1A sequence was aligned to the DnaK sequence to identify regions with loop insertions and deletions. The HspA1A fragment library was produced using the fragment picker. The lowest scoring model was used to produce a model of the complex between HspA1A and the JD of DnaJB8. The structural images were produced using PyMOL.

### Cell biological and biochemical analysis of DnaJB8-Clover cell lines

The human *DNAJB8* protein-coding sequence was cloned using Gibson assembly into a modified FM5 lentiviral expression plasmid^46^ in which the *UbC* promoter was replaced by a *CMV* promoter, the linker sequence was replaced by “GSAGSAAGSGEF” and the *YFP* was replaced by *mClover3*. The resulting gene produced a DNAJB8-mClover3 fusion protein. In parallel, we produced a construct that expresses the fluorescent protein (mClover3) but lacks DnaJB8. Both plasmids we separately co-transfected into HEK293T cells along with helper plasmids (pCMV-VSV-G and psPAX2) to produce lentivirus, which was harvested after 48hrs and used to produce polyclonal cell lines that expressed either DnaJB8-mClover3 or mClover3. For crosslinking experiments, cells from a confluent 10cm^2^ cell culture dish were pelleted and lysed using an insulin syringe in 1xPBS with 1mM DTT, 1mM PMSF, 1x EDTA-free Protease Inhibitor Cocktail (Roche), and 1% Digitonin. After spinning at 1000xg for 10min., the lysate was recovered and incubated with a polyhistidine tagged anti-GFP nanobody (plasmid encoding the nanobody was a kind gift from Dr. Judith Frydman and purified as described previously^24^) for 2.5 hours at 4°C. The lysate was then incubated with 25μL NiNTA beads (Clontech) for 1 hour at 4°C for binding. The beads were washed five times with 300μL 1xPBS. The buffer for each wash was removed after pulse spinning the beads via centrifugation. The beads were preincubated for 5 minutes at 37°C and a final concentration of 57mM ADH and 36mM DMTMM were added to each sample. Following a one-minute incubation with chemical crosslinkers, the reaction was quenched with 1mM ammonium bicarbonate. After another pulse spin to remove the buffer, the beads were resuspended in elution buffer, (8M Urea, 0.5M imidazole, pH 7.5). After a final pulse spin, the supernatant was retained and analyzed by mass spectrometry.

### Cross-Linking Reagents

All crosslinking reagents used are commercially available: ADH (Sigma-Aldrich), and mixed light and deuterated ADH (ADH-h_8_/d_8_) (Creative Molecules). DMTMM (Sigma-Aldrich). For all crosslinking experiments, stock solutions were made of each crosslinking reagent. A 1:1 ADH-d_0_/ADH-d_8_ solution was made at 100mg/mL in 1xPBS pH 7.4 (Sigma-Aldrich). DMTMM (Sigma-Aldrich) was prepared at a 120mg/mL concentration in 1xPBS pH 7.

### Cross-Linking Mass Spectrometry

The *ex-vivo* purified DnaJB8 was dialyzed to remove excess imidazole, and transferred into 1xPBS pH 7.4 buffer. For the full-length DnaJB8 experiments, lyophilized DnaJB8 was resuspended in either 1xPBS(150mM) or 1xPBS(285mM) to a concentration of 100μM. The JD_1-77_ and CTD_170-232_ constructs were purified into 1xPBS buffer, and were prepared for XL-MS experiments at 100μM each. 2μM HspA1A in 1xPBS pH 7.4 buffer and mixed with either 40μM DnaJB8, 40μM JD_1-77_, 40μM DnaJB8ΔCTD and 40μM DnaJB8_F→S_ for XLMS experiments and performed in triplicate. All samples were incubated at 37°C while shaking at 350 rpm for 30 minutes. Final concentrations of 57mM ADH d_o_/d_8_ (Creative Molecules) and 36mM DMTMM (Sigma-Aldrich) were added to the protein samples and incubated at 37°C with shaking at 350 rpm for 30 minutes. The reactions were quenched with 100mM ammonium bicarbonate and incubated at 37°C for 30 min. Samples were lyophilized and resuspended in 8M urea. Samples were reduced with 2.5mM TCEP incubated at 37°C for 30 min followed by alkylation with 5mM iodoacetimide for 30 minutes in the dark. Samples were diluted to 1M urea using a stock of 50mM ammonium bicarbonate and trypsin (Promega) was added at a 1:50 enzyme-to-substrate ratio and incubated overnight at 37°C shaking at 600 rpm. 2% (v/v) formic acid was added to acidify the samples following overnight digestion. All samples were run on reverse phase Sep-Pak tC18 cartridges (Waters) eluted in 50% acetonitrile with 0.1% formic acid. 10μL of the purified peptide fractions was injected for LC-MS/MS analysis on an Eksigent 1D-NanoLC-Ultra HPLC system coupled to a Thermo Orbitrap Fusion Tribrid system. Peptides were separated on self-packed New Objective PicoFrit columns (11cm x 0.075mm I.D.) containing Magic C18 material (Michrom, 3μm particle size, 200Å pore size) at a flow rate of 300nL/min using the following gradient. 0-5min = 5 %B, 5-95min = 5-35 %B, 95-97min = 35-95 %B and 97-107min = 95 %B, where A = (water/acetonitrile/formic acid, 97:3:0.1) and B = (acetonitrile/water/formic acid, 97:3:0.1). The mass spectrometer was operated in data dependent mode by selecting the five most abundant precursor ions (m/z 350-1600, charge state 3+ and above) from a preview scan and subjecting them to collision-induced dissociation (normalized collision energy = 35%, 30ms activation). Fragment ions were detected at low resolution in the linear ion trap. Dynamic exclusion was enabled (repeat count 1, exclusion duration 30sec).

### Analysis of Mass Spectrometry Results

All mass spectrometry experiments were carried out on an orbitrap Fusion Lumos Tribrid instrument available through the UTSW proteomics core facility. Each Thermo .raw file was converted to .mzXML format for analysis using an in-house installation of xQuest^47^. Score thresholds were set through xProphet^47^, which uses a target/decoy model. The search parameters were set as follows. For grouping light and heavy scans (hydrazide crosslinks only): Precursor mass difference for isotope-labeled hydrazides = 8.05021 Da for ADH-d_0_/d_8_; maximum retention time difference for light/heavy pairs = 2.5 min. Maximum number of missed cleavages = 2, peptide length = 5-50 residues, fixed modifications = carbamidomethyl-Cys (mass shift = 57.02146 Da), mass shift of light crosslinker = 138.09055, mass shift of monolinks = 156.1011 Da, MS^1^ tolerance = 15 ppm, and MS^2^ tolerance = 0.2 Da for common ions and 0.3 Da for crosslink ions; search in enumeration mode. For zero-length crosslink search: maximum number of missed cleavages = 2, peptide length = 5-50 residues, fixed modifications carbamidomethyl-Cys (mass shift = 57.02146 Da), mass shift of crosslinker = -18.010595 Da, no monolink mass specified, MS^1^ tolerance = 15 ppm, and MS^2^ tolerance = 0.2 Da for common ions and 0.3 Da for crosslink ions; search in enumeration mode. The FDRs of all *in vitro* experiments range from 0.05 to 0.33.

### Western blot analysis

10μL aliquots of the HEK control, Clover, and DnaJB8-Clover cell-lines were removed from the elution and loaded onto a 4-12% Bis-Tris SDS-PAGE gel for western blotting. Upon running the gel to completion, the gel transferred onto a transfer membrane soaked in Novoblot transfer buffer. Following the transfer, the membrane was soaked in milk blocking buffer for 1 hour at room temperature. For immunolabelling, we added 1:2000 dilution of polyclonal Anti-GFP (rabbit) (Rockland; 600-401-215; 35460) or Anti-DnaJB8 (rabbit) (abcam; ab235546; GR3229943-2) in milk and incubated the membrane shaking at room temperature for 2 hours. The primary antibody solution was dumped and the membrane washed three times for 10 minutes each with 1xTBST before adding the polyclonal Anti-Rabbit IgG Peroxidase (GE Healthcare; NA9340V; 16908235) at a 1:5000 dilution in milk. The membrane was incubated with secondary antibody at room temperature for 1 hour before removing the antibody solution. The membrane was washed 3 times in 5-minute intervals with 1xTBST and finally one 5-minute wash with 1xTBS. The membrane was soaked in 1mL of Luminol enhancer and peroxide solution for 1 minute before imaging.

### In cell analysis of DnaJB8-Clover and Clover cell lines

HEK293T cells were treated with 1x and 3x amounts of lentivirus expressing either DnaJB8-mClover3 or mClover alone were plated at 300,000 cells per well in media (10%FBS, 1%Pen/Strep, 1%GlutaMax in DMEM) in a 6-well glass bottom plate (Cellvis, P06-1.5-N). After 30 hours, cells were stained with Hoescht33342 at a final concentration of 2ug/mL in cell media for 30 minutes at 37C and 5%CO2. The plate was placed on an IN Cell 6000 Analyzer (GE Healthcare) with a heated stage and fifty fields of view were imaged under DAPI and FITC channels at 60X magnification (Nikon 60X/0.95, Plan Apo, Corr Collar 0.11-0.23, CFI/60 lambda). Images were exported as TIFF files for downstream analysis. DnaJB8-mClover3, mClover3, and wild-type HEK293 cells were plated and imaged in triplicates. Total cell counting was done using the CellProfiler v3.0 software^48^ by selecting for DAPI (total cells in acquired images) and mClover3 (total expressing cells in acquired images). Punctae-containing cells were counted manually by two different observers and the data reported as the average with standard deviation between both observers. Expression of Clover and DnaJB8-Clover in 1x and 3x cell lines was quantified from Western blot analysis and by fluorescence intensity of Clover quantified using ImageJ^49^.

### Recombinant expression and purification of DnaJB8 and DnaJB8ΔCTD

The vector used for DnaJB8 expression was a pET-29b vector containing the gene for human DnaJB8, a T7 promoter to activate DnaJB8 expression, a His tag region at the end of the gene, and a gene for kanamycin resistance. The DnaJB8ΔCTD fragment was cloned into pet29b using Gibson assembly. The same protocol was used to express and purify the WT DnaJB8 and DnaJB8ΔCTD proteins. The vector constructs were transformed into *E. coli* BL-21 (DE3) cells and plated onto 2xLB plates containing 0.05mg/mL Kanamycin. 12mL of 2xLB, 0.05mg/mL Kanamycin were prepared and inoculated with a single colony from the plate. This small culture was incubated overnight at 37°C while shaking at 220rpm. In the morning, the 12mL culture was added to 1L of 2xLB supplemented with 0.05mg/mL Kanomycin and incubated at 37°C while shaking at 220rpm. Once OD600 reached 0.6-0.8 A.U., 1mL of 1M IPTG was added to induce DnaJB8 expression. After incubation for an additional 4 hours, the cells were harvested by spinning down the culture at 4,000g for 20 minutes. The resulting cell pellet was resuspended in 25mL 1xPBS and 1mM PMSF in preparation for insoluble fraction separation. The resuspended cells were sonicated at 30% power, 5X pulse for 10 minutes using an Omni Sonic Ruptor 4000 (Omni International). The lysed cells were pelleted at 10,000g for 30 minutes and the supernatant was discarded. The insoluble pellet was rinsed with 1xPBS, 0.75%Tween-20 and again pelleted at 10,000g for 30 minutes.

The insoluble cell pellet was resuspended in 50mL lysis buffer (8M Guanidinium HCl, 50mM HEPES, 20mM Imidazole, 1mM DTT pH 7.5) and sonicated at 30% power, 3X pulse for 1 minute to solubilize the DnaJB8 from the insoluble pellet. After a 30-minute incubation at room temperature, the cellular debris was pelleted at 15,000g for 30 minutes. The resulting supernatant was mixed with 2mL HisPur™Ni-NTA Resin (Thermo Scientific) for 1 hour before being loaded onto a gravity column. The column was washed with an additional 50mL of lysis buffer, followed by 50mL of a second wash buffer (50mM HEPES, 20mM Imidazole, 1mM DTT pH 7.5 in H_2_O). The protein was eluted with 30mL of elution buffer (50mM HEPES, 500mM Imidazole, 1mM DTT pH 7.5 in H_2_O) and collected in 2mL fractions. After selecting for fractions with high purity, the DnaJB8 solution was loaded into 3.5kDa cutoff Biotech CE Dialysis Tubing (Spectrum Labs) and dialyzed overnight at 4°C in 50mM ammonium formate to minimize assembly. The protein was then lyophilized and stored at -80°C for future use.

### Dynamic Light Scattering

All samples were prepared at 1.2mg/mL in 1xPBS, 1mM DTT, pH 7.4. All protein samples were filtered through a 0.22μm PES sterile filter and loaded in triplicate onto a 384 well clear flat-bottom plate. The plate was loaded into a Wyatt DynaPro Plate Reader III and set to run continuously at room temperature at a scanning rate of 1 scan per 15 minutes, with 1 scan composed of 10 acquisitions. The data were analyzed using the Wyatt Dynamics software version 7.8.2.18. Light scattering results were filtered by Sum of Squares (SOS) <20 to eliminate statistical outlier acquisitions within each scan. For DnaJB8 in 1xPBS(150mM) buffer, one of the triplicates contains partial data due to high SOS values. This is a result of increasing polydispersity and heterogeneity, which is consistent with oligomers of that size. R_h_ of observed particles for three time points (0h, 10h, and 20h) were reported as histograms as a function of mass%. The mass% contribution of smaller particles in the full-length DnaJB8 runs in 1xPBS(150mM) and (285mM) buffer were reported as a function of mass% over time using the SOS filter and a size filter of <10nm. Data for DnaJB8_F→S_, JD_1-77_ and CTD_170-232_ was reported as a function of R_h_ over time with the SOS filter applied.

### Modeling of full length DnaJB8 using Rosetta and XL-MS restraints

Given the globular conformations of the JD and CTD we considered how the crosslinks identified for the full-length protein could guide the JD-CTD interaction. Using Rosetta we assembled a monomeric conformation leveraging the JD and CTD conformations while keeping the G/F and S/T regions fully expanded. This starting model was then used in a relax protocol in conjunction with crosslinks (Supplementary Data 1) as constraints to produce an ensemble of 1000 collapsed conformations. A representative low scoring model was selected for further analysis. For acid-acid and acid-lysine contacts 21 and 16 Å distance thresholds were used as restraints. The HYDROPRO^50^ software was used to calculate radii of hydration from structural models.

### Conservation mapping

DnaJB8 homolog sequences were identified using Blast^51-52^ and the sequences were aligned using Clustal Omega^53^. The protein sequence alignment and structure of DnaJB8 JD (PDBID 2DMX) were used as input in Al2Co^54^ to map the conservation onto the structural models. The conservation was mapped onto the models in PyMOL.

### Coevolutionary variation analysis

The GREMLIN software^29-31^ was used to identify covarying amino acid pairs from a DnaJB8 protein sequence alignment. A probability of 0.7 was used to threshold the data to identify amino acids with strong coupling.

### Solid state NMR analysis

Chaperone oligomers were prepared in PBS buffer with different NaCl concentrations. Lyophilized DnaJB8 uniformly labeled with ^13^C and ^15^N (U-^13^C,^15^N) was re-suspended in 1ml of PBS buffer in which the final NaCl concentration was 100mM or 285mM, respectively, for the two samples measured by MAS NMR. Each sample was packed into a 3.2mm MAS NMR rotor (Bruker Biospin) by sedimentation, using a previously described ultracentrifugal packing tool^55^. Sedimentation was done at 175,000 x g force in an Optima L-100 XP Ultracentrifuge using a SW-32 Ti centrifuge rotor for 1 hour. Subsequently, excess of the supernatant fluid was removed. Sample tubes were washed with another 1mL buffer solution after which a second packing step using the same parameters was performed. Finally, the supernatant was removed, spacers were placed on the top of the hydrated sedimented protein oligomers, and rotors were closed with the drive cap, and sealed with a small amount of epoxy to avoid sample dehydration.

Experiments were performed on Bruker 600 MHz and 750 MHz spectrometers, at 277 K temperature using triple-channel (HCN) 3.2mm MAS EFree probes. All experiments were done using TPPM^56^ proton decoupling of 83 kHz during acquisition. Single pulse excitation (SPE) measurements were performed using a 6μs 90° pulse on ^13^C, 3s recycle delay, 1k scans and 1D Cross-Polarization experiments (CP) were performed using a 3.1μs 90° ^1^H pulse, 2ms contact time, recycle delay of 3s, 1k scans. 1D refocused INEPT experiments were done at using 3μs 90° proton pulse, 4μs 90°pulse on ^13^C, recycle delay of 2s and 1k scans. The ^13^C-^13^C 2D CP-DARR experiments were performed using 25ms mixing time, 0.4ms contact time, 3.1μs 90° proton pulse, 6μs 90°pulse on ^13^C, recycle delay of 2.8s, 128 scans. The 2D NCA experiment was performed using 3μs 90° proton pulse, 900μs and 3.25ms ^1^H-^13^C and ^13^C-^15^N contact times respectively, 8μs 180°pulse on ^15^N, recycle delay of 2s, 576 scans. The amino acid type and secondary structure were predicted using the PLUQ program^38^ applied to the chemical shifts in the 2D ^13^C-^13^C spectrum. Linewidth analysis was done using the UCSF Sparky NMR analysis program^57^. Spectral acquisitions were done with Bruker Topspin software and processing was done with the NMRpipe and Sparky software packages ^57-59^.

### Simulations and synthetic NMR spectra

The structure of the DnaJB8 JD in solution was determined previously using solution NMR^42^, allowing us also to generate a synthetic ^13^C-^13^C 2D spectrum using the corresponding solution NMR chemical shifts from the BMRB (entry 11417). To simulate approximate 2D NMR spectra of the other three domains, we made use of the results of MD simulations of full length DnaJB8. The starting DnaJB8 conformation was produced using ROSETTA with a fully expanded conformation of the G/F and S/T domains while keeping the JD and CTD in the folded conformations. MD simulations were prepared using Maestro^60-61^ and carried out in Desmond running the amber99 forcefield. Estimated chemical shifts of the resulting structural models were generated using the SPARTA+ package^39^.

### Recombinant Expression and Purification of DnaJB8_F→S_, J Domain, and C-Terminal Domain

Both vector constructs containing DnaJB8 JD and CTD respectively were cloned into pET-29b using Gibson assembly. The DnaJB8_F→S_ construct was purchased from Genscript and cloned into pET-29b. These vectors were transformed into *E. coli* BL-21 (DE3) cells and plated onto 2xLB plates with 0.05mg/mL Kanamycin. 12mL of 2xLB with 0.05mg/mL Kanamycin were prepared and inoculated with a single colony from each plate. These small cultures were incubated overnight at 37°C while shaking at 220rpm. In the morning, the 12mL culture was added to 1L of 2xLB supplemented with 0.05mg/mL Kanamycin and incubated at 37°C while shaking at 220rpm. Once OD600 reached 0.6-0.8 A.U., 1mL of 1M IPTG was added to induce protein expression. After incubation for an additional 4 hours, the cells were harvested by spinning down the culture at 4,000g for 20 minutes. For preparing ^15^N JD, a single colony was inoculated into 10mL 2xLB supplemented with 0.05mg/mL Kanamycin and incubated for 7-8 hours at 37°C while shaking at 220rpm. The 10mL culture was then mixed into 100mL of M9 minimal media (42mM Na_2_HPO_4_, 22mM KH_2_PO_4_, 8.5mM NaCl, 0.1mM CaCl, 2mM Mg_2_SO_4_, 1E-4% Thiamine, 0.4% Glucose, 187mM NH_4_Cl, 0.05mg/mL Kanamycin) and incubated overnight at 37°C while shaking at 220rpm. In the morning of the following day, the cells were spun down at 2,000g for 10minutes and resuspended in 20mLs of M9 minimal media containing 15N labeled NH_4_Cl in place of the unlabeled molecule. This was immediately added to 1L of M9 minimal media with ^15^N labeled NH_4_Cl and allowed to incubate at 37°C shaking at 220rpm. Once OD600 reached 0.6-0.8 A.U., 1mL of 1M IPTG was added to induce protein expression. After incubation for an additional 4 hours, the cells were harvested by spinning down the culture at 4,000g for 20 minutes. The cell pellets were resuspended in 20mL of soluble wash buffer (SWB) (50mM KPO_4_, 300mM NaCl, 10% glycerol, 1mM PMSF, 10mM BME, pH 8) and sonicated at 30% power, 5X pulse for 10 minutes using an Omni Sonic Ruptor 4000 (Omni International). After incubation at room temperature for 1 hour, the cell lysate was spun down at 15,000g for 30 minutes to separate the soluble supernatant from the insoluble pellet. The supernatant was mixed with 2mL TALON® Metal Affinity Resin (Clontech) and incubated shaking at 4°C for 1 hour. The protein-resin slurry was loaded onto a gravity column and washed with an additional 40mL of SWB. This was followed by subsequent washes: 20mL SWB with 0.5% tritonX-100, 20mL SWB with adjusted 700mM NaCl, 20mL SWB with 0.1mM ATP and 5mM MgCl_2_, and an additional 40mL of SWB. The protein was eluted with 16mL SWB with 200mM Imidazole into 2mL fractions. After selecting for fractions with high purity, the protein solution was loaded into 3.5kDa cutoff Biotech CE Dialysis Tubing (Spectrum Labs) and dialyzed overnight at 4°C in 1xPBS to restore native folding. Both domain constructs were further enriched by running on a GE Superdex 200 Increase 10/300 column in 1xPBS 1mM DTT pH 7. The protein was aliquoted and flash frozen in liquid nitrogen and stored at -80°C for future use.

### SEC-MALS

DnaJB8_F→S_, JD_1-77_, and CTD_170-232_ constructs at a concentration of 5.1 mg/mL, 5.0 mg/mL, and 6.6 mg/mL, respectively, in 1xPBS were filtered through a 0.1 μm filter to remove larger impurities. Each sample was further filtered using a 0.22 μm centrifugal filter before 100 μL was applied to a Superdex 200 Increase 10/300 column equilibrated in 1xPBS. The column was in line with a Shimadzu UV detector, a Wyatt TREOS II light-scattering detector, and a Wyatt Optilab tREX differential-refractive-index detector. The flow rate was 0.5 mL/min. The data were analyzed with Wyatt’s ASTRA software version 7.1.0.29. SEDFIT^62^ was used to calculate the dn/dc of the protein.

### Circular Dichroism

Recombinant CTD_170-232_ domain constructs were transferred into 10mM NaPO_4_, 150mM NaF, pH 7.4 buffer at a concentration of 40μM. The experiment was run using a Jasco J-815 Circular Dichroism instrument with a PMT detector using a 10 mm quartz cuvette. 6 accumulations were taken at a speed of 50 nm/min along the UV spectrum from 190 nm to 300 nm. Spectra analysis was done using the BeStSel online software^63-64^ to determine secondary structural composition.

### CTD model generation

Fragment libraries for the CTD sequence were generated using the Robetta server. 5000 models were produced using the *ab initio* protocol and clustered to identify unique conformations. Lowest scoring models from the top clusters showed high structural similarity. The identified crosslinks from the CTD in full length DnaJB8 or isolated CTD were evaluated for consistency with each model in the ensemble. For acid-acid and acid-lysine crosslinks a distance threshold of 21 and 16 Å, respectively, was considered as satisfied and consistent with the chemistry. Distances were calculated using a custom script in MATLAB ver.R2019a and the crosslink pairs were visualized using PyMOL. The HYDROPRO^50^ software was used to calculate radii of hydration from structural models.

### Fluorescence polarization

JD_1-77_ was labelled with 10x FITC-maleimide (sigma) in 1xPBS 1mM TCEP for 2 hours at room temperature. The reaction was quenched and the excess dye was removed using a Zeba spin desalting column (Thermo). For all experiments, 0.2μM JD_1-77_ was incubated in triplicate with a titration gradient of Hsp70 (150μM-0μM) or CTD_170-232_(150μM-0μM) in 1xPBS, 1mM TCEP, pH 7.4. For competition experiments, labelled JD_1-77_ was mixed with 3.125μM CTD in triplicate and incubated at room temperature for 1 hour before adding a titration gradient of Hsp70(150μM-0μM). Fluorescence polarization readings were taken with excitation at 494nm and emission at 525nm. The data were fit to a one site specific binding model using GraphPad Prism 7.04.

### Microscale Thermophoresis

The MST binding experiments were performed on Nanotemper Monolith NT.115 in the Molecular Biophysics Resource core at UTSW and analyzed with a standard protocol^65^. Binding measurements were performed in triplicate. The JD_1-77_ was labeled with FITC-maleimide (sigma) and 200nM JD_1-77_-FITC was titrated by a serial 2-fold dilution of CTD_180-232_. Data were fit in PALMIST^65^ to a 1:1 binding model.

### Solution NMR with ^15^N-labeled JD and CTD

The 300μM ^15^N-labeled JD was exchanged into 20mM Tris 100mM NaCl 1mM DTT pH 7 buffer in preparation for solution NMR. Each HSQC run was done for 4 hours at 1 scans/min with the temperature fixed at 299K. After each run, unlabeled CTD was titrated into the sample at 1:1, 1:3, and 1:6 ratios sequentially. All scans were collected on an Agilent DD2 600MHz instrument at the UT Southwestern Biomolecular NMR Facility. Each spectrum was converted into a readable format and phase corrected using NMRPipe^58^. Peak assignments were based on the deposited information from BMRB (11417). The software Sparky^57, 59^ was used to analyze the peak shifts across all spectra.

### Recombinant expression and purification of HspA1A

HspA1A gene was cloned into the pMCSG7 plasmid^66^ and transformed into BL-21 (DE3) cells and plated onto 2xLB plates with 0.1mg/mL ampicillin. 12mL of 2xLB with 0.1mg/mL ampicillin were prepared and inoculated with a single colony from each plate. These small cultures were incubated overnight at 37°C while shaking at 220rpm. In the morning, the 12mL culture was added to 1L of 2xLB supplemented with 0.1mg/mL ampicillin and incubated at 37°C while shaking at 220rpm. Once OD600 reached 0.6-0.8 A.U., 1mL of 1M IPTG was added to induce protein expression. The cells continued to incubate overnight at 12°C shaking at 220rpm. After incubation, the cells were lysed using a PandaPlus 2000 homogenizer (GEA) by pressing the cells with 10,000p.p.m pressure. The lysate was spun at 15,000xg for 45 minutes to remove insoluble cell components, and the resulting supernatant was mixed with 2mL TALON® Metal Affinity Resin (Clontech) and incubated at 4°C for 1 hour. The slurry was spun down at 700xg for 2 min to remove the majority of the buffer and the beads were added onto a gravity column. The beads were washed with 6CV of wash buffer (50mM Tris, 500mM NaCl, 10mM imidazole, 5mM βME, pH 8) and eluted with 5mL of elution buffer (50mM Tris, 500mM NaCl, 300mM imidazole, 5mM βME, pH 8). HspA1A containing fractions were confirmed by SDS-PAGE and pooled together for desalting. Desalting/buffer exchange was performed using a PD-10 desalting column (GE Healthcare), where HspA1A fractions were transferred into anion-exchange wash buffer (50mM Tris, 20mM NaCl, 1mM DTT, pH 8.75). The protein was loaded onto a HiTrap Q HP anion exchange column (GE Healthcare) and eluted across a gradient of anion-exchange elution buffer (50mM Tris, 1M NaCl, 1mM DTT, pH 8.75). HspA1A-containing fractions were once again combined and loaded onto a Superdex™ 200 Increase 10/300 GL (GE Life Sciences) size exclusion column, where HspA1A was further purified and transferred into 1xPBS, 1mM DTT, pH 7.4 buffer for all subsequent experiments.

## Acknowledgements

This work was supported by grants from the Welch Foundation and the Effie Marie Cain Endowed Scholarship (L.A.J.) and NIGMS R01 GM112678 (P.V.D.W.). We appreciate the help of the Molecular Biophysics Resource core, Structural Biology Laboratory, Biomolecular Nuclear Magnetic Resonance Facility, and Proteomics Core Facility at the University of Texas Southwestern Medical Center. The 750 MHz ssNMR instrument at the University of Pittsburgh was acquired with funding from NIH grant 10 OD012213-01.

## Author Contributions

B.D.R., I.M., P.V.D.W., and L.A.J. conceived and designed the overall study. B.D.R. performed *in vitro* protein binding assays, cell models, crosslink mass spectrometry, and ROSETTA simulations. I.M. and P.V.D.W. performed ssNMR experiments and analyzed the data. S.B. performed DnaJB8 CTD experiments and ROSETTA simulations. J.V.A produced mammalian cell lines and collected microscopy images. B.D.R., I.M., P.V.D.W., and L.A.J. wrote the manuscript, and all authors contributed to its improvement.

## Data availability

The data sets generated during and/or analysed during the current study are available from the corresponding authors on reasonable request.

## SUPPLEMENTARY INFORMATION

### Supplementary Data

**Supplementary Data 1. Primary XL-MS data for DnaJB8 and its domains**.

**Supplementary Data 2. Primary DLS measurements for DnaJB8 oligomers**.

### Supplemental Tables

**Supplementary Table 1.** Residue distribution per domain with net charge for each domain.

| Domain<br>aa.<br>length | Domain | Residue |  |  |  |  |  |  |  |  |  |  |  |  |  |  |  |  |  |  |  | Net charge<br>for the<br>domain |
| --- | --- | --- | --- | --- | --- | --- | --- | --- | --- | --- | --- | --- | --- | --- | --- | --- | --- | --- | --- | --- | --- | --- |
|  |  | A | R | N | D | Q | E | G | H | I | L | K | M | F | P | S | T | Y | V | W | C |  |
| 1-75 | JD | 8 | 5 | 3 | 6 | 1 | 7 | 2 | 1 | 1 | 6 | 10 | 1 | 1 | 3 | 7 | 0 | 5 | 4 | 2 | 1 | +2 |
| 76-148 | G/F | 6 | 5 | 2 | 5 | 0 | 5 | 13 | 2 | 1 | 2 | 0 | 1 | 14 | 6 | 6 | 3 | 2 | 0 | 1 | 0 | -5 |
| 149-185 | S/T | 0 | 0 | 1 | 0 | 0 | 1 | 7 | 1 | 0 | 1 | 1 | 3 | 4 | 0 | 15 | 4 | 0 | 1 | 0 | 1 | 0 |
| 186-232 | CTD | 0 | 2 | 3 | 2 | 3 | 6 | 4 | 7 | 2 | 2 | 6 | 1 | 0 | 0 | 2 | 3 | 0 | 6 | 1 | 0 | 0 |

**Supplementary Figure 1.**
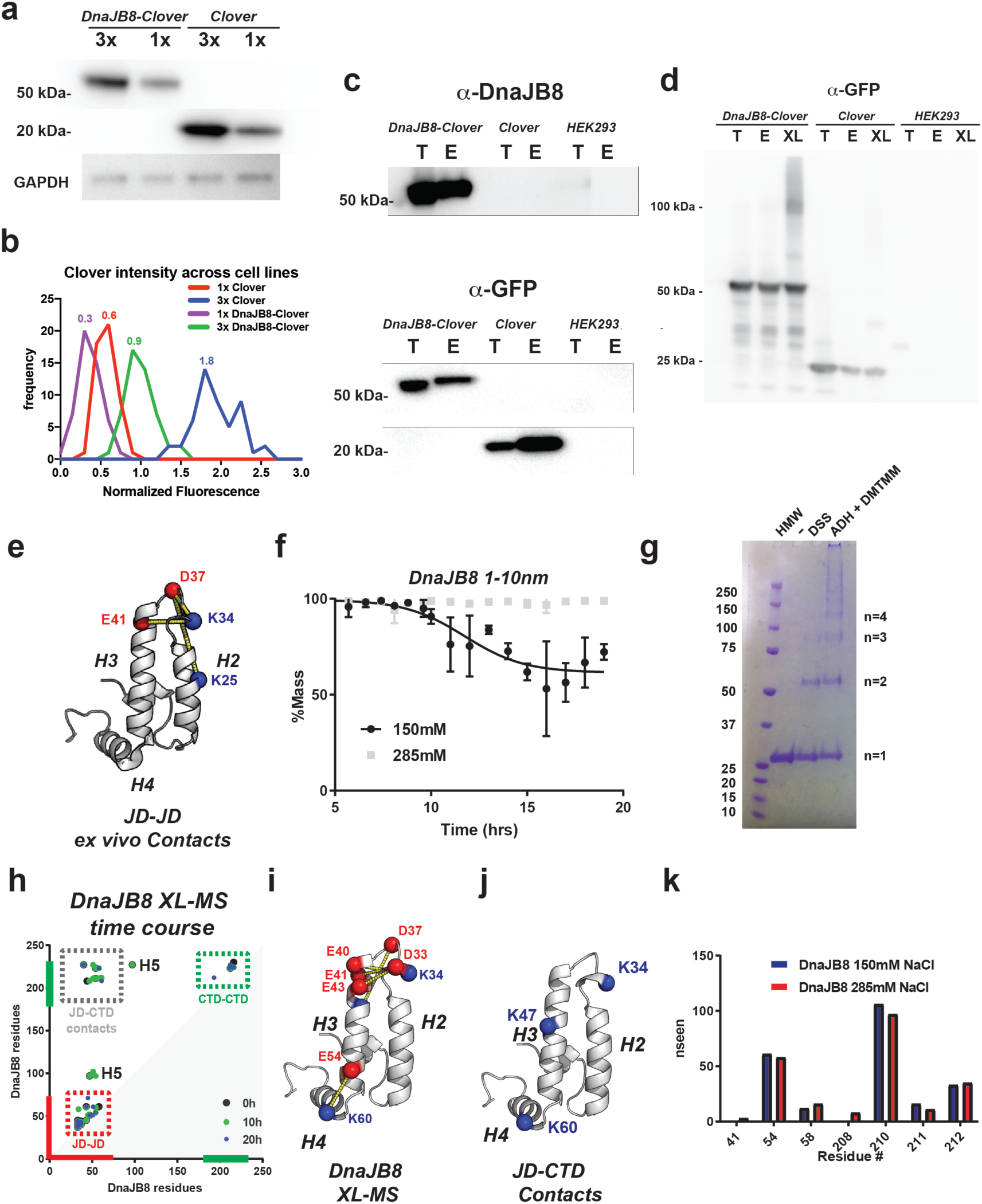
Biochemical and mass spectrometry analysis of DnaJB8 in cells and *in vitro*. **(a)** Western blot analysis of 1x DnaJB8-Clover, 3x DnaJB8-Clover, 1x Clover and 3x Clover cell lines to quantify levels of expression in each cell line. GAPDH western blot is shown as a loading control. (**b**) Normalized FITC intensity analysis of 1x Clover (red), 3x Clover (blue), 1x DnaJB8-Clover (purple) and 3x DnaJB8-Clover (green) cell lines. Intensity measurements were calculated from 50 images for each cell line and the signal normalized to DAPI fluorescence intensity. (**c**) Western blot analysis of input and anti-GFP nanobody elutions DnaJB8-Clover, Clover and HEK293 cell lines as detected with anti-DnaJB8 and anti-GFP antibodies. Total (T) and elutions (E) for each condition are shown. (**d**) Western blot analysis of DnaJB8-Clover and Clover proteins isolated from mammalian cell lines. Total (T), elutions (E) and crosslinked elutions (XL) of each condition are shown. Western blot was probed with GFP antibodies. **(e)** JD intra-domain ADH/DMTMM crosslinks identified from DnaJB8-Clover isolated from cells mapped onto the JD structure. JD is shown in cartoon representation and is colored in white. Sites of crosslink are shown as spheres and are colored red or blue for aspartic/glutamic and lysines, respectively. Dashed yellow lines connect linked amino acid pairs. **(f)** DnaJB8 particles with a R_h_ of 1-10nm in the DLS data were analyzed by proportion of the total sample (% mass) and binned by the size distribution of the constituent particles. The time evolution of this mass fraction is shown for 150mM NaCl (black) and 285mM NaCl buffer conditions in 1xPBS. **(g)** SDS-PAGE coomassie gel showing DnaJB8 in the absence of crosslinker (left), with DSS (middle), and with DMTMM and ADH (right). **(h)** XL-MS contact map of DnaJB8 crosslinks identified using DMTMM and ADH from a time course: t=0hrs (large black dot), t=10hrs (medium green dot) and t=20hrs (small blue dot), small. The axes are colored in red and green for JD and CTD, respectively. Crosslink pairs between JD-CTD are shown in dashed box colored grey, red and green, respectively. Helix 5 crosslinks are denoted by H5. **(i)** JD intra-domain ADH/DMTMM *in vitro* crosslinks mapped onto the JD structure. JD is shown in cartoon representation and is colored in white. Sites of crosslink are shown as spheres and are colored red or blue for aspartic/glutamic and lysines, respectively. Dashed yellow lines connect linked amino acid pairs. **(j)** Three *in* vitro JD-CTD inter-domain ADH/DMTMM crosslinks mapped onto the JD structure. DnaJB8 JD is shown in cartoon representation and is colored in white. JD lysine sites that crosslink to CTD are shown as spheres and are colored blue. **(k)** Histogram of frequency of ADH monolinks observed in normal and elevated ionic strength XLMS experiments.

**Supplementary Figure 2.**
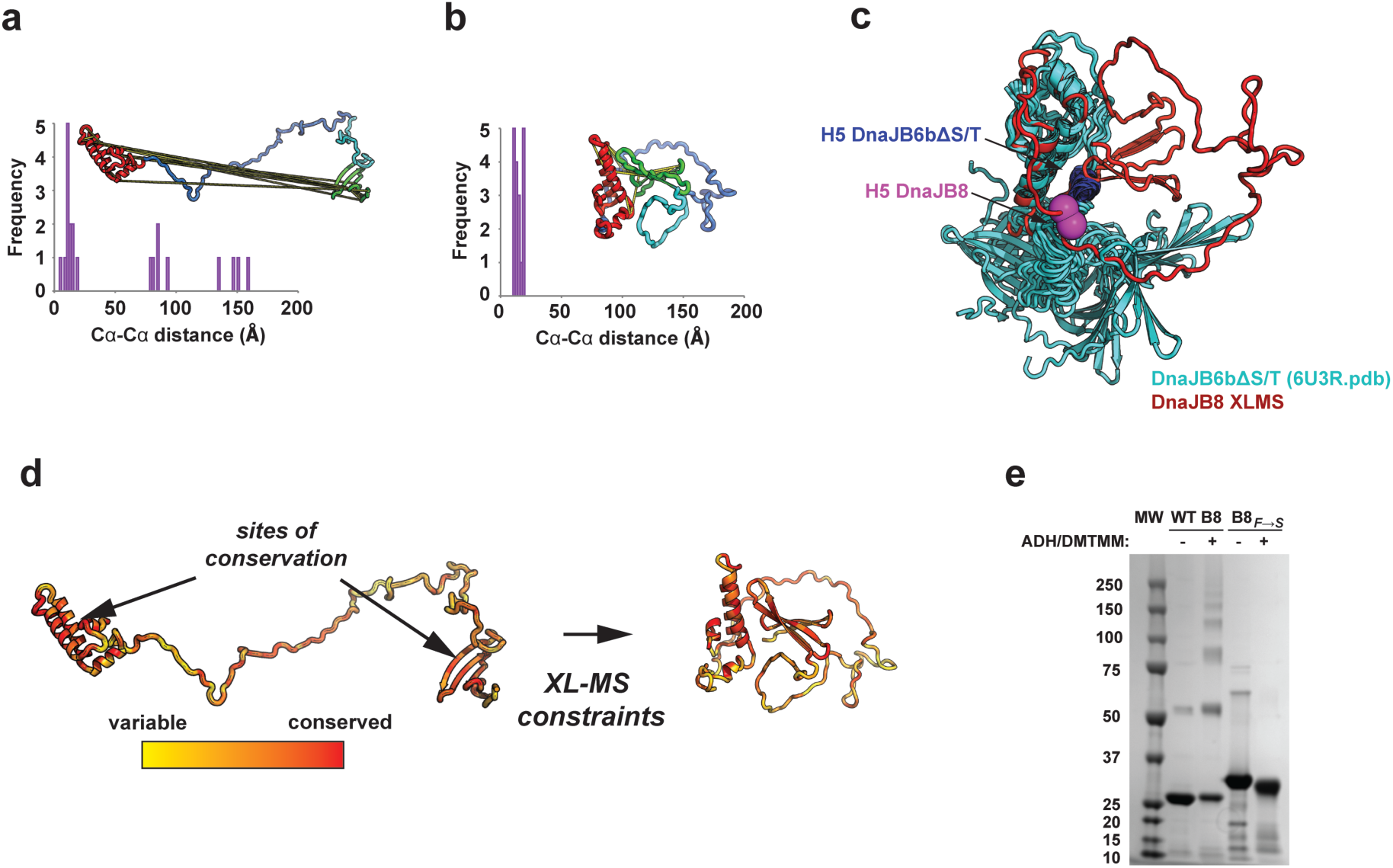
Modeling the full length DnaJB8 monomer using crosslinks. **(a)** DnaJB8 ADH/DMTMM crosslinks mapped onto an expanded structure featuring the conserved JD and our predicted CTD model. The histogram of C_α_-C_α_ distances shows 11 crosslinks that exceed the physical limitations for ADH/DMTMM. **(b)** Rosetta *ab initio* collapsed structure of DnaJB8 with ADH/DMTMM crosslinks mapped. The histogram of C_α_-C_α_ distances shows that all crosslinks are satisfied when the JD and CTD are docked in this model. **(c)** Overlay of our Rosetta *ab initio* collapsed structure (red) with the solution NMR ensemble of the published DnaJB6bΔST deletion variant (cyan; PDBID: 6U3R) The proposed helix 5 (H5) is shown on the DnaJB8 (pink) and DnaJB6bΔST (blue) models. The position of H5 in our DnaJB8 XL-MS-constrained model is close to the position in the DnaJB6b model. The major difference is the position of the CTD, which is influenced by both the XL-MS constraints, and the inclusion of the S/T region that was deleted in the DnaJB6b construct. **(d)** Conserved surfaces on the JD and CTD in the full-length DnaJB8 model mediate the interaction. Highly conserved sites are colored in red (and highlighted by arrows) and variable positions are colored in yellow. Sequence-based conservation of DnaJB8 shows highly conserved faces along the JD, and some conserved faces in the CTD that overlap with the XL-MS identified surfaces. **(e)** SDS-PAGE coomassie gel of DnaJB8 and DnaJB8_F→S_ mutant crosslinked with ADH/DMTMM.

**Supplementary Figure 3.**
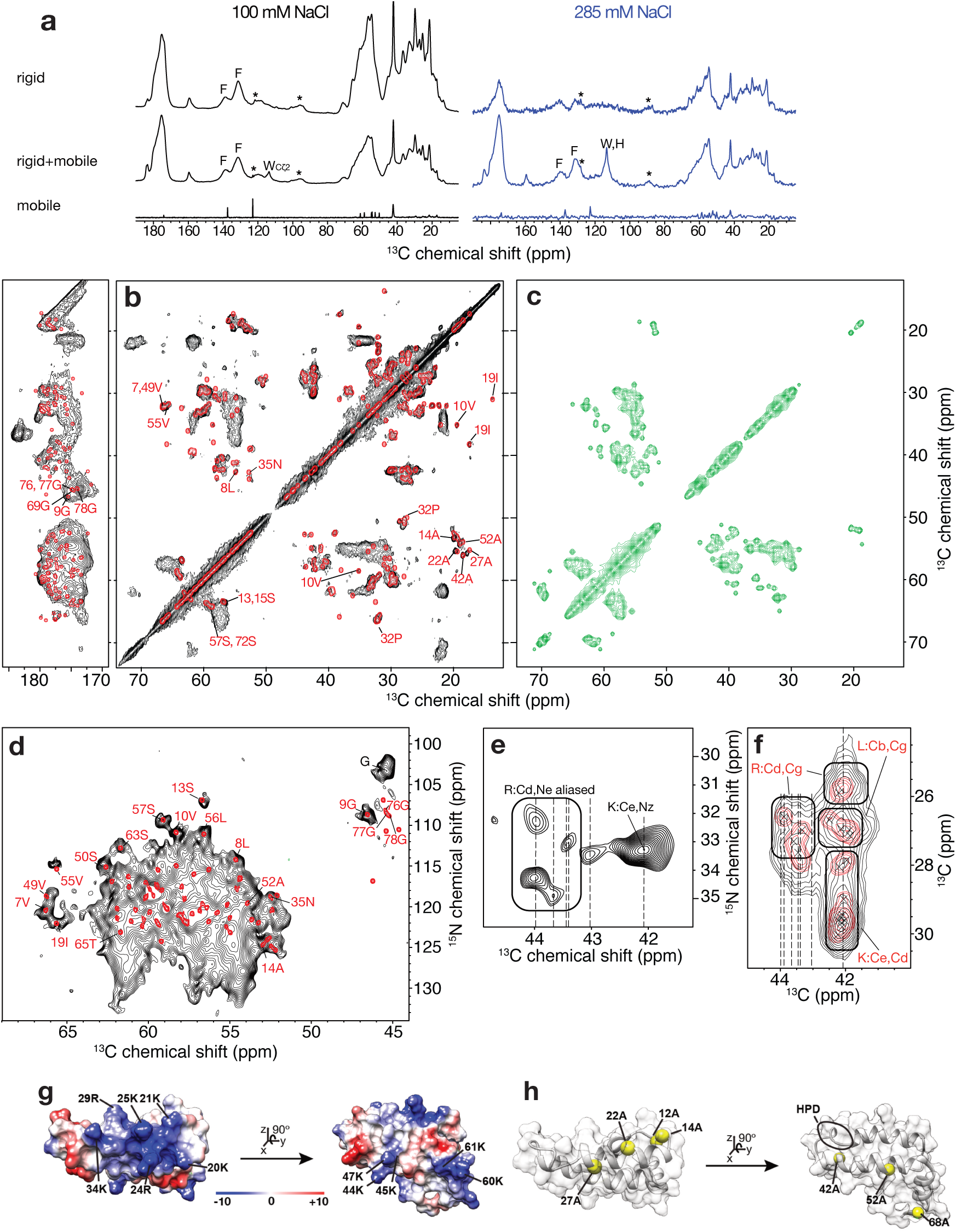
Additional ssNMR data and analysis for DnaJB8 oligomers in PBS. **(a)** ^13^C 1D spectra at 100mM NaCl (black) and elevated ionic strength (blue) that show rigid residues (CP), rigid and mobile residues (SPE) and only mobile residues (INEPT). **(b)** Overlay of the experimental (black) 2D ^13^C-^13^C CP-DARR spectrum of DnaJB8 oligomers uniformly labelled with ^13^C and ^15^N and synthetic (red) spectrum of the DnaJB8 JD in solution generated from BMRB chemical shifts (red assignments). **(c)** Simulated spectrum from three other domains (G/F, S/T and CTD), showing diagonal peaks and one-bond cross-peaks involving backbone Cα-Cβ and CO-Cα carbons. **(d)** 2D ^13^C-^15^N CP-based NCA experimental spectrum (black) of DnaJB8 oligomers uniformly labeled with ^13^C and ^15^N, overlaid with synthetic spectrum of the DnaJB8 JD in solution (red) from BMRB chemical shifts (red assignments). **(e)** 2D CP-based ^13^C-^15^N spectrum with Cε-Nζ and (aliased) Cδ-Nε correlations from immobilized Lysine and Arginine side chains, respectively. **(f)** Enlarged ^13^C-C spectral region from panel (A) demonstrating good alignment of Arg, Lys and Leu signals between experimental ssNMR (black) and solution chemical shifts of the JD (red). Dashed lines mark known ^13^C positions used to confirm amino acid types of signals. **(g)** Electrostatic surface potential of the DnaJB8 JD, with selected residues indicated. Highly positive and negative potentials are colored blue and red, respectively. White color represents neutral charge. **(h)** The same view showing the location of Ala with narrow peaks at 100 mM NaCl which disappear at 285 mM NaCl (Fig. 3, panel “c”).

**Supplementary Figure 4.**
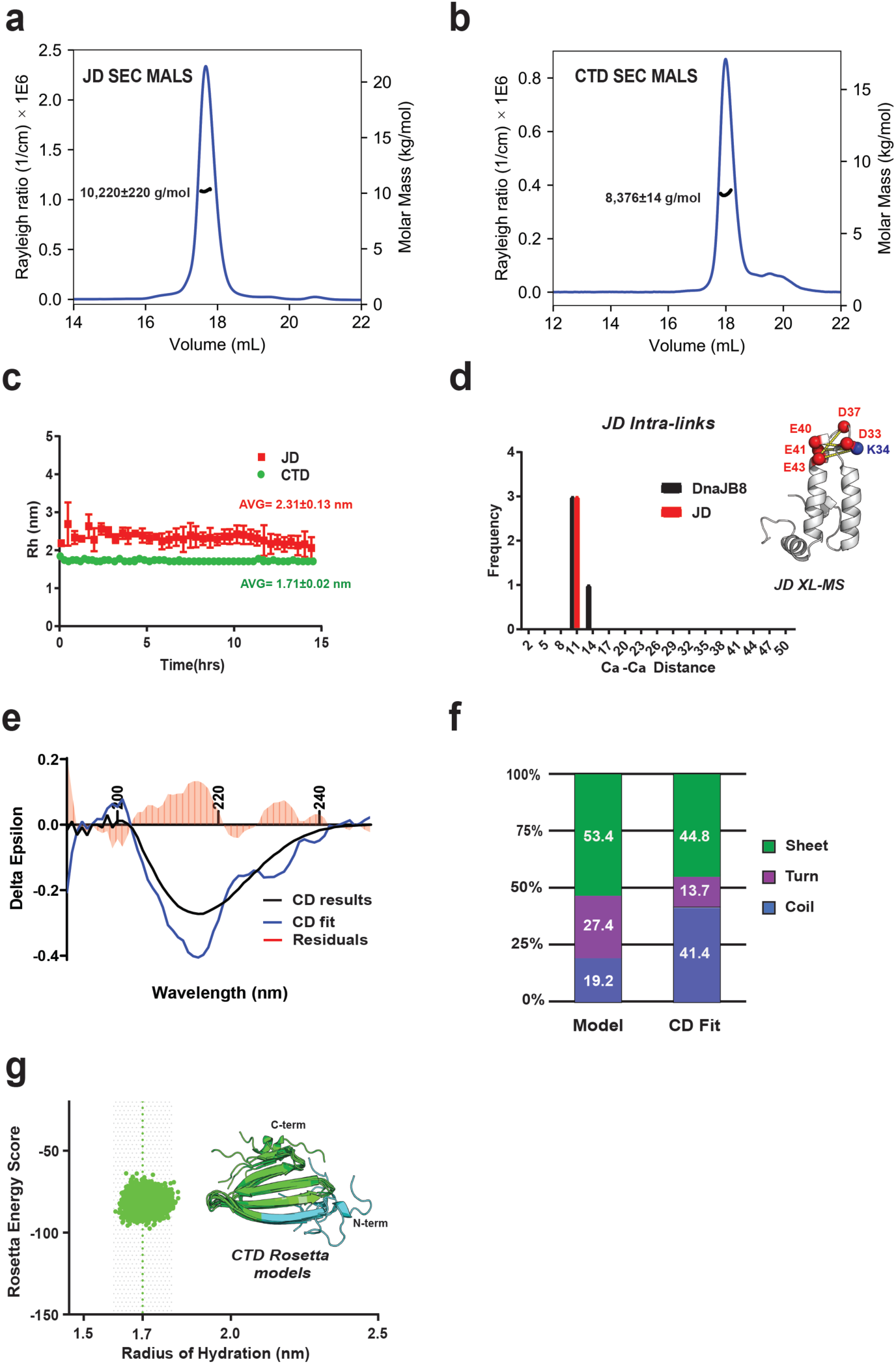
JD and CTD are stable monomers. **(a)** SEC-MALS of JD_1-77_ shows a single peak with a calculated molar mass of 10,220 ± 220 g/mol consistent with a monomer. **(b)** SEC-MALS of CTD_170-232_ shows a single peak with a calculated molar mass of 8,376 ±14 g/mol consistent with a monomer. **(c)** DLS time course of JD_1-77_ and CTD_170-232_ constructs. The average R_h_ of JD_1-77_ and CTD_170-232_ was calculated to be 2.30±0.12 nm and 1.71±0.02 nm, respectively. **(d)** Histogram of JD intra-domain crosslinks across XLMS experiments for DnaJB8(black), JD_1-77_(red), and JD_1-77_ mixed with CTD_170-232_(grey). All intra-domain crosslinks satisfy the physical constraint of <20nm for the ADH/DMTMM crosslinkers. **(e**,**f)** CD spectra of CTD was analyzed using the Bestsel server. Secondary structure analysis of the data shows high β-sheet character (yellow) with some helix character (purple) and no random coil (blue). The results are compared to the CTD model generated from ab initio. **(g)** 5,000 Rosetta *ab initio* generated models of CTD. The structural models are consistent with R_h_ values derived from DLS (dashed line). Structural overlay of low energy scoring models reveals consistent pleated beta-sheet fold. Models are shown in cartoon representation and are colored in green.

**Supplementary Figure 5.**
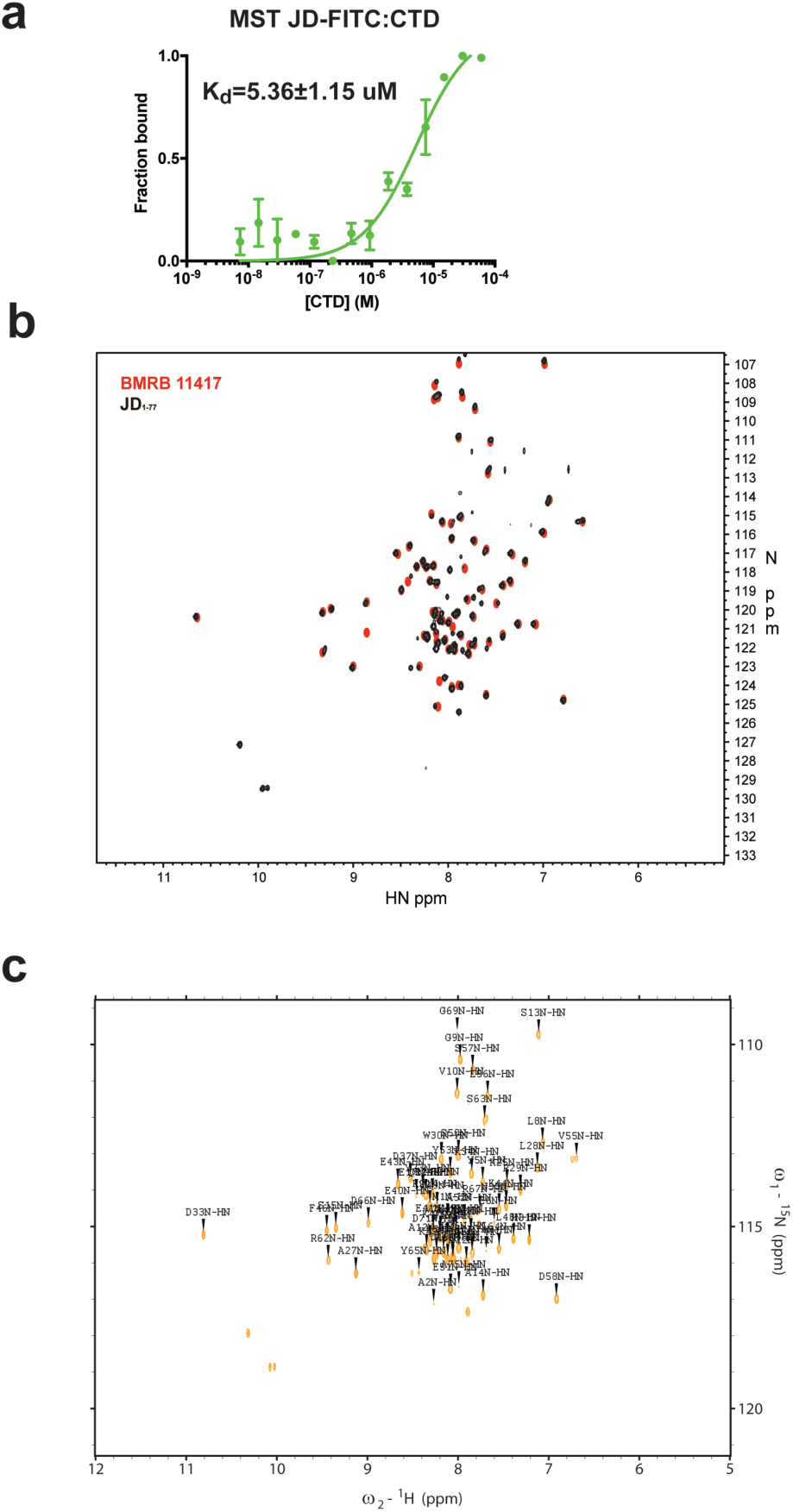
JD_1-77_ construct spectra consistent with deposited chemical shifts. **(a)** Microscale Thermophoresis measurements of JD_1-77_-FITC binding to CTD_170-232_. Measurements were performed in triplicate using 200nm JD_1-77_-FITC and a range of CTD_170-232_. The samples were incubated and measured in a Monolith NT.115 instrument. The data were processed using PALMIST software and the normalized fluorescence values plotted as an average across three replicates with standard deviations **(b)** Overlay of our ^15^N-^1^H HSQC spectra for JD_1-77_(black) onto the submitted spectra for DnaJB8 JD (red) (BMRB:11417). 97% of assigned peaks in the published spectra were matched to our data. (**c)** Amino acid assignments mapped onto the HSQC spectra for the JD_1-77_ protein.

**Supplementary Figure 6.**
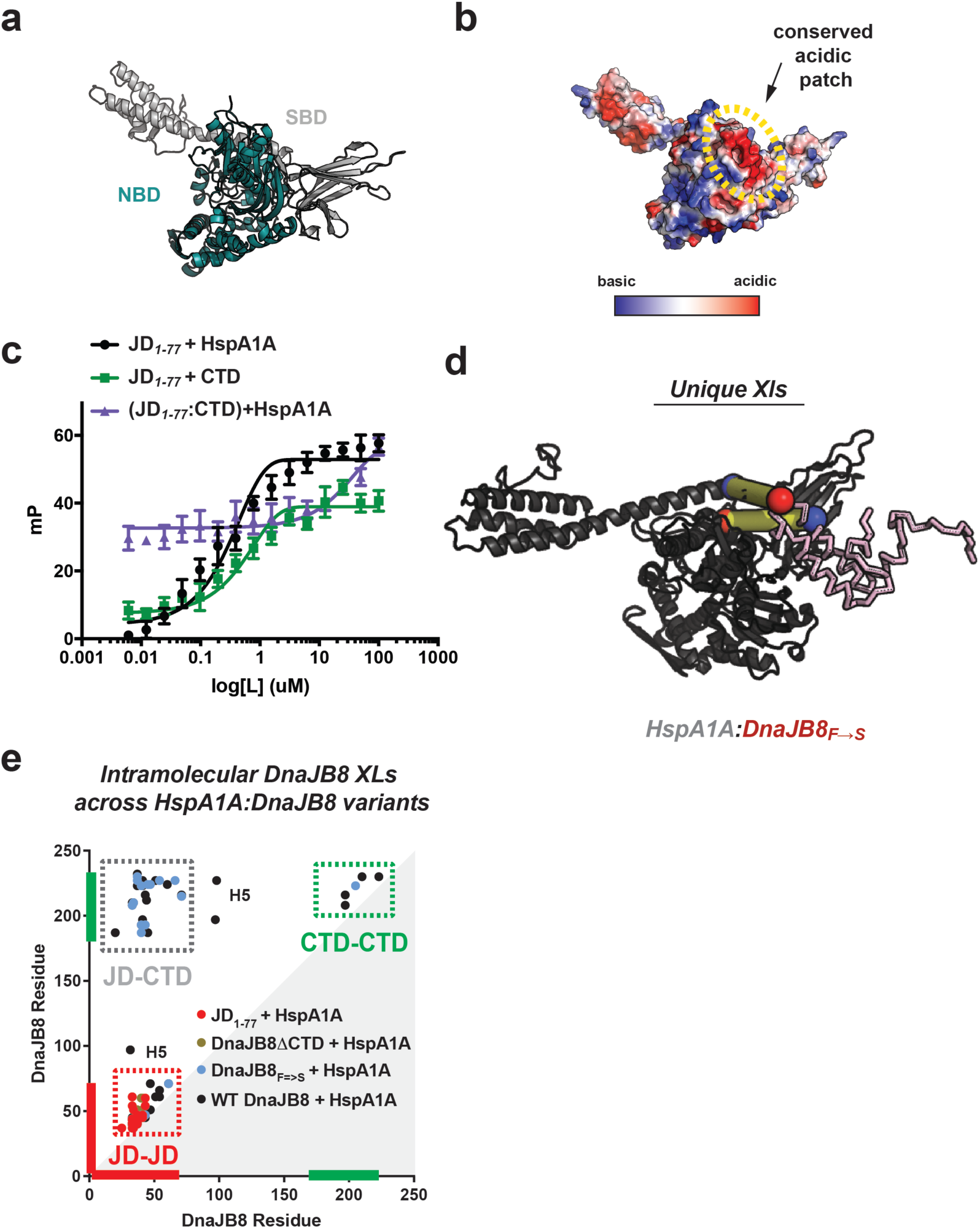
Hsp70 proteins present a conserved acidic surface. **(a)** Structure of HspA1A colored by domain with the nucleotide binding domain (NBD) in turquoise and substrate binding domain (SBD) in grey. (**b)** Electrostatic surface mapped onto the HspA1A structure with the negatively charged Hsp40-JD-binding site circled in yellow. **(c)** Raw FP binding curves of labeled JD mixed with HspA1A (black), with CTD (green) and (JD-CTD)+HspA1A (purple). Y-axis is defined as milipolarization (mP). FP experiments were performed in triplicate and shown as averages with standard deviation. **(d)** The two intermolecular crosslinks identified across three XL-MS datasets between JD and HspA1A in the DnaJB8:HspA1A complex mapped onto JD-HspA1A model. JD is shown in ribbon representation and HspA1A in cartoon representation, colored pink and black respectively. Sites of crosslink are shown as red or blue spheres for aspartic/glutamic acid and lysines, respectively. Yellow lines connect linked amino acid pairs. **(e)** XL-MS contact map of changes in intramolecular DnaJB8 crosslinks identified using DMTMM and ADH in co-mixtures of different variants of DnaJB8 with HspA1A: WT DnaJB8:HspA1A (black), DnaJB8ΔCTD:HspA1A (brown), DnaJB8_F→S_:HspA1A (blue) and JD_1-77_:HspA1A (red). The axes are colored in red and green for JD and CTD, respectively. Crosslink pairs between JD-CTD are shown in dashed box colored grey, red and green, respectively. Contacts to helix 5 in WT DnaJB8 are denoted with H5.

